# Optics-free Spatial Genomics for Mapping Mouse Brain Aging

**DOI:** 10.1101/2024.08.06.606712

**Authors:** Abdulraouf Abdulraouf, Weirong Jiang, Zihan Xu, Zehao Zhang, Samuel Isakov, Tanvir Raihan, Wei Zhou, Junyue Cao

## Abstract

Spatial transcriptomics has revolutionized our understanding of cellular network dynamics in aging and disease by enabling the mapping of molecular and cellular organization across various anatomical locations. Despite these advances, current methods face significant challenges in throughput and cost, limiting their utility for comprehensive studies. To address these limitations, we introduce *IRISeq* (Imaging <u>R</u>econstruction using Indexed <u>Seq</u>uencing), a optics-free spatial transcriptomics platform that eliminates the need for predefined capture arrays or extensive imaging, allowing for the rapid and cost-effective processing of multiple tissue sections simultaneously. Its capacity to reconstruct images based solely on sequencing local DNA interactions allows for profiling of tissues without size constraints and across varied resolutions. Applying *IRISeq*, we examined gene expression and cellular dynamics in thirty brain regions of both adult and aged mice, uncovering region-specific changes in gene expression associated with aging. Further cell type-centric analysis further identified age-related cell subtypes and intricate changes in cell interactions that are distinct to certain spatial niches, emphasizing the unique aspects of aging in different brain regions. The affordability and simplicity of *IRISeq* position it as a versatile tool for mapping region-specific gene expression and cellular interactions across various biological systems.

**One Sentence Summary:** *IRISeq*, an innovative optics-free spatial transcriptomics method, uncovers aging-related changes in spatial gene expression and focal cell interactions in brain aging.

## Main Text

Understanding the spatial organization of molecules and cells is crucial for studying cellular network dynamics in aging and disease. For example, recent advancements in spatial transcriptomics, by multiplexed *in situ* hybridization(*1–3*) or indexed oligonucleotide capture arrays(*4–10*), have revolutionized our ability to spatially profile genome-wide RNA expression across diverse anatomical locations. These technologies offer detailed insights into cellular networks, significantly enhancing our understanding of the cellular interactions essential for maintaining organismal functions.

Despite these advances, current spatial transcriptomic methods face significant challenges in throughput and cost, which hinder their ability to quantify the alterations in cellular interactions across multiple regions, conditions, and independent replicates. For example, while multiplexed *in situ* hybridization provides subcellular resolution(*1–3*), it is limited by imaging speed and requires specialized equipment and pre-selected gene probes. On the other hand, oligonucleotide capture arrays face limitations in their throughput and the ability to cover extensive areas: *prior* indexing methods (*e.g., 10X Genomics Visium* (*4*)*, DBiT-seq*(*6*)), use predefined DNA barcodes at specific positions, whereas *posterior* indexing methods, such as *slide-seq* (*5*, *11*), *HDST* (*7*) and *Stereo-seq*(*8*), distribute unidentified DNA barcodes on a 2D surface and later identify them through imaging techniques like sequencing. Scaling up these indexing approaches is technically challenging, constrained by issues such as barcode pre-assignment or the reliance on multiple rounds of optical imaging.

The approach of “spatial mapping by sequencing” offers a promising solution to these challenges by utilizing barcoded DNA sequences to record the spatial proximity of millions of biological molecules(*12*). This strategy, previously employed in DNA Hi-C for constructing three-dimensional chromatin structures(*13*), is recently being adapted to profile spatial interactions of several targeted mRNA molecules for inferring cellular spatial interactions in cultured cell lines (*e.g., DNA microscope*(*14*)) or the spatial distribution of cellular surface proteins(*15*). Despite these innovations and the development of supporting theoretical frameworks(*14*, *16*, *17*), we still lack a robust, high-throughput, and cost-effective “spatial mapping by sequencing” platform for region-specific transcriptomic analysis of complex tissues.

Here, we present *IRISeq* (Imaging <u>R</u>econstruction using Indexed <u>Seq</u>uencing), a scalable and cost-effective spatial genomic analysis method that relies exclusively on sequencing, eliminating the need for predefined capture arrays or optical imaging. This approach utilizes DNA barcoded beads to capture gene expression across each of thousands of spatial locations, with the global spatial positions of these beads decoded through their interaction signals with adjacent beads. This method allows for the parallel processing of multiple tissue sections in a single day at a significantly lower cost than commercial platforms and without specialized equipment. Furthermore, *IRISeq* offers the flexibility to map genome-wide RNA expression across a range of resolutions (from 5 µm to 50 µm) and capture areas (from 0.6 cm to 1.5 cm in diameter). Meanwhile, we developed computational pipelines designed to reconstruct tissue images solely from sequencing data. This pipeline also facilitates the integration of single-cell and spatial data for assessing region-specific cellular distributions and interactions. We applied *IRISeq* to analyze brain sections from both adult and aged mice and revealed widespread and region-specific gene expression changes, as well as alterations in spatial cell-cell interactions associated with brain aging. The resulting spatially resolved atlas provides a detailed view of molecular and cellular changes in the aging brain, demonstrating the applicability of *IRISeq* for exploring spatial gene expression dynamics in complex biological tissues.

## Overview of IRISeq (Imaging <u>R</u>econstruction using Indexed <u>Seq</u>uencing)

The optimized *IRISeq* protocol comprises several key steps (**Fig. 1A**, **fig. S1**): (i) Bead preparation: two types of oligo-barcoded beads are prepared: ’receiver beads’ coated with PolyT sequences to capture nearby cellular mRNA, and ’sender beads’ with a photocleavable linker and a PolyA sequence. These barcoded beads are created through a split-pool ligation approach (*18*), such that each bead has its unique barcode (**fig. S1A**). (ii) Photocleavage and oligo capture: beads are evenly distributed on a glass slide. A UV device is then utilized to photocleave oligos from the sender beads, which then diffuse and are captured by the receiver beads, mimicking the capture of tissue mRNA (**fig. S1B**). The bead array is then frozen on dry ice to stabilize it. (iii) Tissue transfer and mRNA capture: frozen tissue sections are transferred onto this array using cryosectioning. The mRNA from the tissue is then captured by the receiver beads through hybridization and tissue digestion processes. (iv) Reverse transcription and sequencing: post tissue digestion, beads are collected for reverse transcription and PCR, followed by sequencing to obtain transcriptome data and bead connection details. Detailed *IRISeq* protocols are provided in the supplementary files to enable cost-efficient spatial transcriptomic mapping of large tissue sections in individual laboratories (**Supplementary Note 1**, **Table S1**).

**Figure 1:**
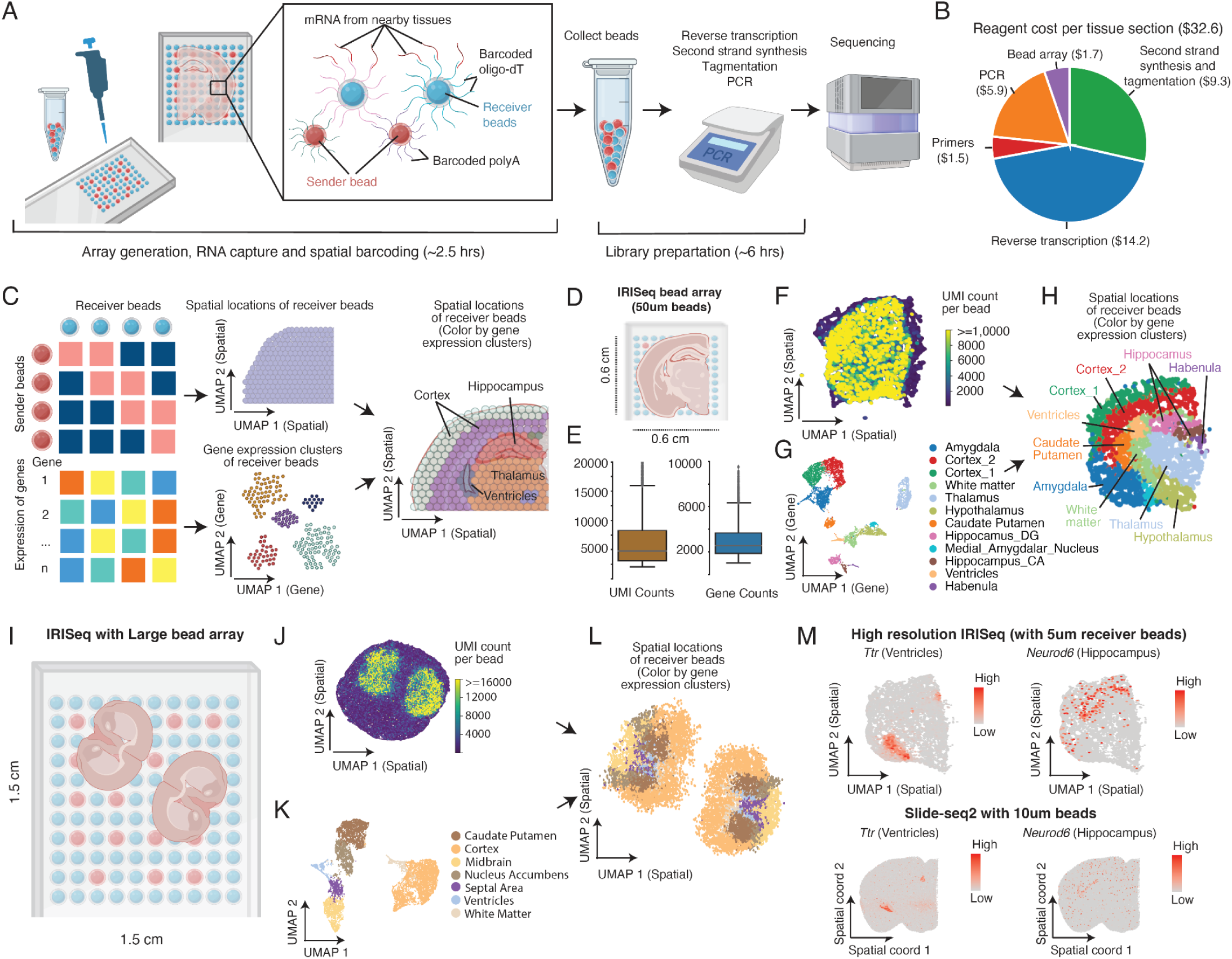
Overview and Validation of the *IRISeq* Platform. (**A**) Schematic illustrating the *IRISeq* experimental pipeline utilizing barcoded gel beads (‘receiver beads’) to capture mRNA expression from nearby cells, along with barcoded oligos from adjacent ‘sender beads’ for spatial localization. (**B**) Pie chart detailing the reagent cost breakdown for profiling a tissue section using a 0.6 cm x 0.6 cm bead array. (**C**) Diagram showing the generation of gene count and bead-bead connection matrices from *IRISeq* data to identify region-specific gene expression and infer bead spatial locations for image reconstruction. (**D**) Depiction of a small-scale *IRISeq* experiment using a 0.6 cm x 0.6 cm bead array to profile a mouse brain hemisection. (**E**) Box plots displaying the distribution of unique transcripts and genes detected per receiver bead in the small-scale experiment. (**F**) UMAP plot showing the spatial distribution of receiver beads based on interactions with sender beads, colored by the number of unique transcripts per bead. (**G**) We integrated the gene expression data of receiver beads with spatial transcriptome data from *10x Visium*(*20*). The UMAP plot shows the integrated gene expression clusters, with each bead colored by annotated brain regions. (**H**) UMAP plot visualizing the spatial distribution of receiver beads, with each bead colored by annotated brain regions. (**I**) Depiction of a large-area *IRISeq* experiment using a 1.5 cm x 1.5 cm bead array to profile two entire brain sections. (**J**) UMAP plot of receiver beads from the large-area *IRISeq* experiment, colored by the number of unique transcripts detected per bead. (**K-L**) UMAP plots displaying gene expression clusters (K) and reconstructed spatial distributions (L) of receiver beads from the large-area experiment, colored by annotated brain regions. (**M**) UMAP plots comparing the spatial distribution of 5 µm receiver beads in *IRISeq* (Top) with beads from published Slide-seqV2 data(*21*) (Bottom), colored by expression of region-specific markers *Ttr* and *Neurod6*. *IRISeq* and Slide-seq2 data are normalized and scaled per bead (**Methods**).

After library preparation and sequencing, the output from *IRISeq* includes a bead-bead interaction counts matrix, which identifies connections between sender and receiver beads, and a gene expression matrix that details transcripts mapped to each receiver bead (**Fig. 1C**, left). To determine the global spatial positions of the receiver beads, we applied principal component analysis (PCA) to the bead-bead interaction matrix, followed by Uniform Manifold Approximation and Projection (UMAP) analysis (*19*) (**Methods**, **fig. S2**). The resulting 2D UMAP coordinates will reflect the designed layout of the assay, preserving the positional relationships among the beads (**Fig. 1C**, middle). Furthermore, we utilized dimension reduction and clustering techniques on the gene expression matrix to annotate each receiver bead based on region-specific gene expression patterns (**Fig. 1C**, middle). This was followed by cell-type-specific deconvolution analysis, integrating single-cell RNA-seq data to map diverse cell types and quantify their spatial interactions (**Methods**).

As an initial demonstration of *IRISeq*, we profiled a coronal mouse brain section using a 0.6 cm x 0.6 cm bead array with a bead diameter of approximately 50 μm (**Fig. 1D**). This experiment yielded 5,028 receiver beads, each capturing a median of 4,587 unique transcripts (2,535 genes) and 1,998 unique bead-bead connection oligos, with each receiver bead connecting to a median of five sender beads (**Fig. 1E-F**, **fig. S3A**). Clustering analysis based on the gene expression profiles of all receiver beads identified 12 transcriptionally distinct brain region clusters, including cortical, caudal putamen, amygdalar, ventricles, white matter, and midbrain regions, validated using region-specific gene markers (**Fig. 1G**, **fig. S3B**). Further validation was achieved by integrating the *IRISeq* data with the published *10x Visium* spatial transcriptome dataset (*20*), which showed consistent region-specific gene expression patterns across spatial locations (**fig. S3C-D**). We then mapped the spatial locations of all receiver beads based solely on their connection signals with sender beads (**Fig. 1F**). The resulting reconstructed image (**Fig. 1H**) preserved the local neighborhood structure of the receiver beads and accurately positioned beads from different regions to their expected locations (**fig. S3E-F**), akin to the pre-indexed *10x Visium* oligo array. This success highlights the capability of *IRISeq* to map the spatially barcoded transcriptome without the need for optical imaging.

A distinctive feature of *IRISeq* is its ability to construct the spatial distribution of micro-beads based solely on their local interactions, bypassing the need for pre-indexed bead arrays or imaging to decode transcript locations. This enables the spatial profiling of large tissue sections beyond the typical size limits of conventional methods, such as the 0.65 cm x 0.65 cm restriction with the commercial *10x Visium* platform(*4*). Demonstrating this capability, we constructed a large 1.5 cm x 1.5 cm bead array without subdivisions, allowing the simultaneous profiling of two entire mouse brain coronal sections (**Fig. 1I**). This setup recovered a total of 14,756 receiver beads, each yielding a median of 5,329 unique transcripts and 652 unique bead-to-bead connection oligos. Similar to its performance on the small array, *IRISeq* effectively mapped transcriptionally distinct brain regions to their precise spatial locations on the large array (**Fig. 1J-L**), which underscores its efficacy in spatial transcriptomics analysis for large tissue areas.

Another unique aspect of *IRISeq* is its adaptable resolution, achieved by adjusting the size of the beads used in constructing the bead array, similar to other bead-based spatial transcriptomic methods(*5*, *11*). To illustrate this capability, we constructed a high-resolution bead array using 5µm magnetic receiver beads. This modification allowed for high-resolution spatial transcriptome mapping of a mouse coronal hemisection (**fig. S3G**), resulting in a collection of 27,732 receiver beads, each with a median of 77 unique transcripts, and a range of 26 to 2,620 unique transcripts per bead, comparable to published *Slide-seqV2* (bead size of 10µm) (*11*) (**fig. S3H**). Despite the reduced bead size, the connection signals between beads enabled the precise reconstruction of the global distribution of most sender beads and detailed mapping of gene expression features with high specificities, such as *Ttr* in the ventricle and *Neurod6* in the hippocampus, aligning with data from published spatial transcriptomics(*21*) (**Fig. 1M**). This demonstration confirms the versatility of *IRISeq* for spatially mapping molecular distributions at various resolutions and scales.

In summary, the *IRISeq* platform distinguishes itself from traditional spatial genomic methods through several unique features: (i) Cost-effectiveness: *IRISeq* is significantly more affordable than commercial platforms, costing approximately $30 per tissue section (or less than $1 per mm²) (**Fig. 1B**), compared to over $1,000 per section typically required by other methods(*22*). (ii) Scalability: the ability to process multiple tissue sections in a large-area array facilitates comparative studies of various conditions and replicates within the same experiment. (iii) Optics-free spatial mapping: By relying on local neighborhood interactions for spatial reconstruction, *IRISeq* is particularly well-suited for profiling large tissues, such as human brains, without the need for imaging. (iv) Adjustable resolution: *IRISeq* offers flexibility to explore cellular interactions at different scales by varying bead sizes. This adaptability makes *IRISeq* a versatile tool suitable for a wide array of research applications.

## A spatially resolved transcriptome atlas of the mouse brain

With the *IRISeq* platform, we next conducted a spatial transcriptome analysis of the brains of adult (4-month) and aged (23-month) wild-type C57BL/6 female mice, with three replicates per age group (**Table S2)**. We profiled twenty-four coronal sections in total, including one section each for the frontal and middle part of the cortex (“Section 1” and “Section 2”), and two adjacent sections near the dorsal part of the cortex (“Section 3”) (**Fig. 2A**). These sections covered brain areas from the frontal part of the isocortex to the dorsal aspect of the cortex, including the hippocampus, thalamus, hypothalamus, and other associated brain regions (**Fig. 2A**). Following data processing and quality control to exclude low-quality receiver beads, we recovered a median of 6,358 receiver beads per section, resulting in a total of 128,405 spatially distinct transcriptome profiles (**Fig. 2B**). Of note, these samples were profiled using an earlier version of *IRISeq* (0.6cm x 0.6cm bead array with a bead diameter of ∼50um), which has slightly lower efficiency compared to the optimized version (protocol details in **Supplementary Note 1**). Nevertheless, we successfully recovered an average of 3,682 unique transcripts (median of 2,662) and 6,771 unique bead-bead connection oligos (median of 4,209) per receiver bead (**Fig. 2C-D**).

**Figure 2:**
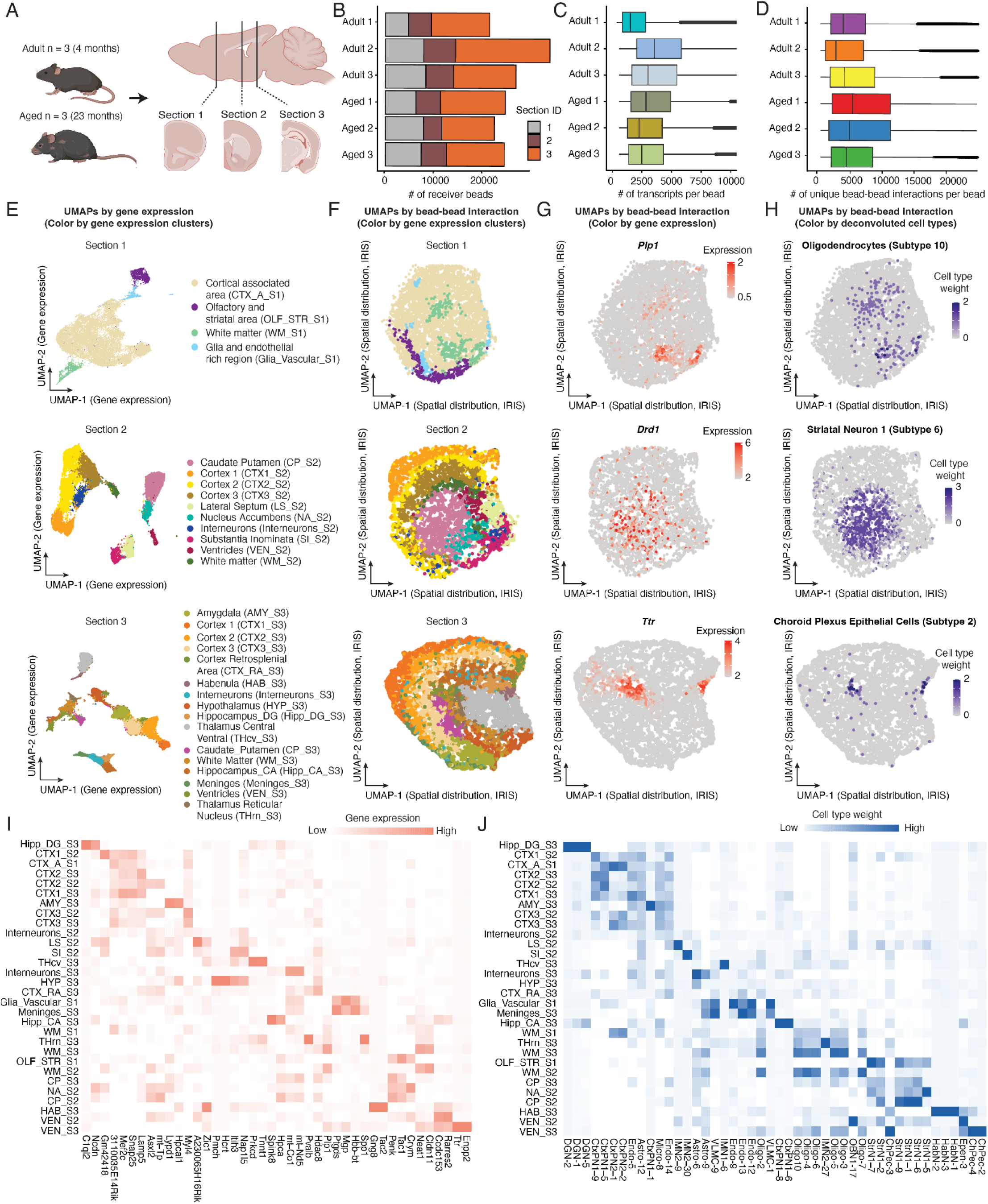
Spatial transcriptomic analysis of mouse brain aging using *IRISeq*. (**A**) Schematic of *IRISeq* profiling across various brain regions, including the frontal isocortex, dorsal cortex, hippocampus, thalamus, hypothalamus, and additional associated regions. “Section 1” and “Section 2” are for the frontal and middle part of the cortex and “Section 3” is near the dorsal part of the cortex. (**B**) Barplot showing the number of spatially barcoded receiver beads recovered per mouse individual. (**C-D**) Boxplots showing the number of unique transcripts (C) and unique bead-bead interactions (D) detected per receiver bead, aggregated across all brain sections for each mouse individual. (**E**) UMAP plots showing the gene expression clusters of receiver beads integrating *IRISeq* data with published spatial transcriptomics datasets (*21*, *24*), colored by annotated brain regions. (**F**) Image reconstruction of all receiver beads derived from bead-bead connections, with coloring corresponding to the annotated regions based on gene expression shown in (E). (**G-H**) Similar to (F), but each bead is colored according to the expression of region-specific gene markers (G) or cell type weight (H) representing cell type abundance by *RTCD* analysis (*25*). (**I**) Heatmap showing the gene expression specificity across regions, with expression data aggregated, normalized, and scaled for each region. (**J**) Cell type distribution across regions, showing sub-cluster IDs from the previous single-cell study (*20*). Cell type weight is calculated using *RCTD*(*25*), integrating published single-cell data(*20*) with *IRISeq* data, then regional subtype proportions are aggregated, normalized, and scaled. Cell type abbreviations: DGN: Dentate Gyrus Neurons. CtxPN1: Cortical Projection Neurons 1. Endo: Endothelial Cells. Astro: Astrocytes. IMN: Interbrain and Midbrain Neurons. VLMC: Vascular Leptomeningeal Cells. Oligo: Oligodendrocytes. OBN: Olfactory Bulb Neurons. StrN: Striatal Neurons. HabN: Habenula Neurons. Epen: Ependymal Cells. ChPec: Choroid Plexus Epithelial Cells. Micro: Microglia.

To annotate spatially barcoded transcriptome profiles across brain regions, we utilized *LIGER*(*23*), a non-negative matrix factorization technique, to integrate gene expression data from *IRISeq* with published spatial transcriptomics datasets profiling corresponding sections (*21*, *24*) (**Methods**). The spatially resolved transcriptomes from *IRISeq* showed a high degree of overlap with published datasets, enabling the identification of thirty distinct brain regions annotated from the existing studies (**Fig. 2E**, **fig. S4A**). Based on bead connection data, we reconstructed the spatial distribution of each receiver bead across different sectioning regions. This reconstruction accurately mapped the receiver beads to specific brain regions, consistent with their gene expression-based annotations and with the published spatial transcriptome datasets (**Fig. 2F**, **fig. S4B**). The region-specific gene expression signatures, such as *Plp1* for white matter, *Drd1* for the caudate putamen, and *Ttr* for the ventricles, further validated the accuracy of the spatial reconstruction (**Fig. 2G**). These results underscore the capacity of the *IRISeq* platform to recover region-specific molecular signatures without reliance on pre-indexed oligo arrays or optical imaging.

To further investigate the spatial distribution of brain cell types, we utilized *RCTD* (*25*), a supervised learning approach that integrates spatial expression data with single-cell transcriptome data for mapping region-specific distribution of heterogeneous cell types. *RCTD* computes mean gene expression profiles for each cell type from the scRNA-seq dataset and generates a spatial cell type map by representing each spatial transcriptomics pixel as a linear combination of these cell types. Using *RCTD*, we integrated the spatial transcriptomics data from *IRISeq* with our previously published brain single-cell transcriptome dataset that has identified over 300 distinct cell subtypes across different age stages(*20*). This integration effectively mapped known brain cell types and subtypes across various regions, such as oligodendrocyte subtypes in white matter, striatal neurons in the caudate putamen, and choroid plexus epithelial cells in the ventricles (**Fig. 2H**). As further validation, we observed a high consistency in cell-type-specific locations inferred by *IRISeq* compared to a spatial transcriptomics dataset generated using the *10X Visium* platform (*20*) (**fig. S5**).

Through differential analysis, we identified highly region-specific gene features and cell types (**Fig. 2I-J, Table S3-4**). For instance, genes like *Ccdc153* and *Rarres2* were highly enriched in the ventricular areas of sections 2 and 3, consistent with previous findings (*26*, *27*). However, section-specific features were also evident, such as the enriched expression of *Enpp2* only in section 3. Similarly, cell type distributions varied by section; OB neuroblast (OBN1-17) enrichment only appeared in section 2, corresponding with its prevalence in the subventricular zone(*28*). This analysis highlights the necessity of sampling multiple brain regions to accurately map the spatial distribution of heterogeneous cell types.

## Age-associated changes in region-specific gene expression programs

We next investigated aging-associated gene expression changes by conducting differentially expressed gene (DEG) analysis across thirty brain regions, identifying 274 genes upregulated and 245 downregulated (FDR of 0.05 with more than three-fold changes between aged and young brains) (**Fig. 3A**, **Table S5**). These changes showed distinct region-specific patterns (**Fig. 3B-C**, **fig. S6A**). For instance, *Sult1c1*, associated with detoxification, hormone regulation, and inflammation(*29*), showed more than a fivefold increase in the aged ventricular region of section 3. *Clec18a*, known for its roles in metabolic and immune responses(*30*), was uniquely upregulated in the aged habenular region. In contrast, the aged hypothalamic region displayed a significant decrease in *Hcrt* expression, which is linked to changes in sleep-wake cycles and appetite control in aging(*31*). The observed age-related, region-specific gene expression changes were further validated using the *10x Visium* platform(*20*) (**fig. S6B**).

**Figure 3:**
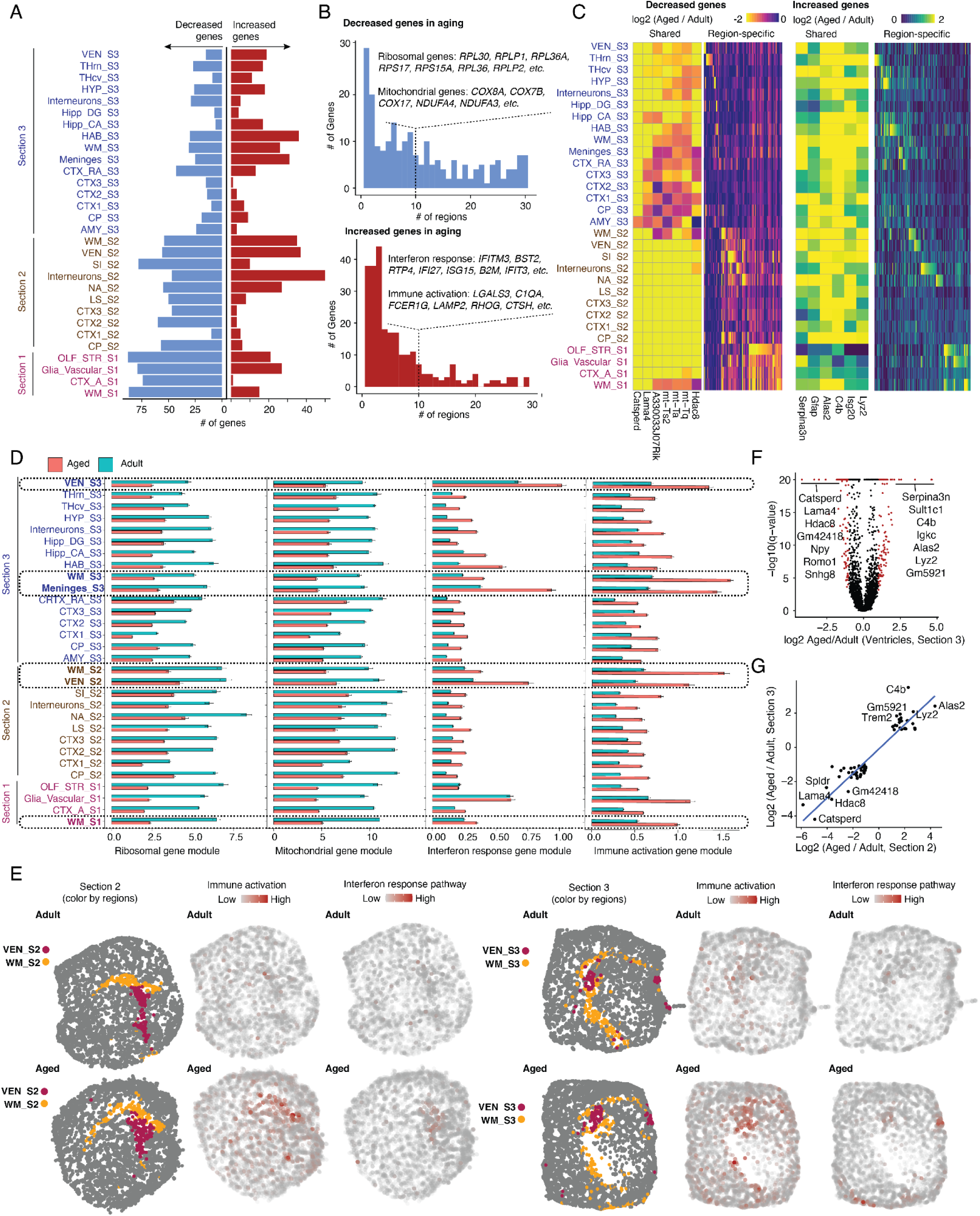
Age-associated changes in region-specific gene expression programs. (**A**) Bar charts depict the number of genes differentially expressed (DE) between adult and aged animals across various brain regions, categorized as upregulated (red) and downregulated (blue). DE genes are defined by more than three-fold changes between the ages with an FDR-adjusted p-value < 0.05. (**B**) Barplot displays the distribution of down-regulated (top) and up-regulated (bottom) DE genes across regions, highlighting genes consistently altered in more than ten brain regions, grouped by pathway. (**C**) Heatmap showing the log2-transformed fold changes in normalized gene expression between aged and adult animals for DE genes across different brain regions, distinguishing shared changes (left) from region-specific changes (right). (**D**) Barplots showing the scaled expression and standard error of gene modules related to ribosomal genes, mitochondrial function, the interferon response, and immune activation across brain regions in both adult (blue) and aged (red) conditions. Gene inclusion for each module is detailed in panel (B). Expression values for each pathway were calculated by aggregating and normalizing pathway-related gene expressions, followed by log transformation and scaling. (**E**) Reconstructed spatial maps display the expression levels of gene modules associated with the complement and interferon response pathways in adults (top) and aged animals (bottom) for Sections 2 (left) and 3 (right). (**F**) Volcano Plot showing differentially expressed genes in the ventricles of Section 3 when comparing aged versus adult samples, with significant changes highlighted in red. (**G**) Scatter plots compare the alterations in differentially expressed genes between ventricle regions from Sections 2 and 3, including a linear regression line. Only DE genes that are identified in both sections are shown.

Notably, we identified 92 downregulated and 47 upregulated genes exhibiting consistent aging-associated changes across more than ten brain regions. Genes showing global downregulation were primarily associated with mitochondrial function (*e.g., mt-Ta, mt-Ts2, mt-Tf, mt-Tq*), oxidative phosphorylation (*e.g., Ndufa1, Ndufa3, Cox8a, Cox6c*), and ribosomal functions (*e.g., Rpl30, Rps15a, Rps17*) (**Fig. 3B**-**D**). These findings are consistent with previous studies (*32*, *33*), indicating significant declines in energy production, cellular metabolism, and protein synthesis in aged brains. Additionally, alterations in genes related to neuronal modulation and signaling (*e.g., Ntsl, Npy, Cck, Tac2*) suggest potential age-related disruptions in neuromodulatory cells, particularly in neuronally enriched regions such as the cortex and the diverse neuromodulatory area of the hypothalamus, which is further confirmed by the published *10x Visium* dataset (*20*) (**fig. S7A-B**).

The 47 genes upregulated across various aged brain regions were significantly enriched in immune response pathways, including the complement pathway (*e.g., C4b, C1qa*), interferon response (*e.g., Isg20, Ifi27, Ifit3*), and antigen presentation (*e.g., Psmb8, H2-Q7, B2m, H2-K1*) (**Fig. 3B-D**). This upregulation, consistent with previous studies (*32*, *34*), indicates increased neuroinflammation with age. These immune responses were particularly pronounced in the ventricles, white matter, and regions abundant in meningeal, endothelial, and ependymal cells (**Fig. 3D**-**E**), indicating the crucial roles of the blood-brain and blood-CSF barriers in neuroimmune dysfunction associated with aging. Additionally, there was a significant upregulation of genes related to tissue structural integrity (*e.g., Col6a1, Col6a2, Hapln2*), suggesting age-related extracellular and intracellular structural remodeling that could affect tissue integrity and cellular function (**fig. S7A-B**).

Focusing on the ventricles, a primary site of inflammation in brain aging (*32*), we identified significant alterations in gene expression between young and aged brains (**Fig. 3F**). The down-regulated genes are predominantly involved in protein synthesis (*e.g., Rpl35a, Rpl34, Rps28*), mitochondrial function (*e.g., Cox8a, Cox7b*), and neurogenesis (*e.g., Hdac8, Stau1, Romo1*). For example, *Hdac8*, linked to DNA stress response, impacts neural stem cell proliferation in the subventricular zone (*35*), while *Stau1* plays a critical role in neural progenitor cell renewal by regulating mRNA stability (*36*). Conversely, up-regulated genes highlight an increase in immune response and inflammation (*e.g., C1qa, C1qb, Ctss, Ifit3*), antigen presentation (*e.g., B2m, H2-D1*), glial activation (*e.g., Gfap*), and non-coding RNA expression (*e.g., Neat1*). These changes suggest a state of chronic inflammation and heightened immune activity in the ventricles, associated with neurodegenerative processes in the aging brain(*37*). Remarkably, the gene expression changes were highly consistent across different ventricular regions from two coronal sections, demonstrating the robustness of our platform in identifying region-specific vulnerable genes in aging (**Fig. 3G**).

## Age-associated changes in the spatial distribution and interaction of different cell types

Using the cell-type-specific deconvolution approach *RTCD*(*25*) as previously described, we integrated *IRISeq* spatial expression data with a published single-cell transcriptome dataset (*20*). This integration enabled us to map previously annotated brain cell types and subtypes across various brain regions, identifying both widespread and region-specific cell types (Figure 2J). We also conducted differential abundance analysis to explore region-specific cell population dynamics during aging. Additionally, by analyzing cell-cell co-locations on the same bead, we quantified cell-cell interactions and examined changes in these interactions across regions (**Methods**, Figure 4A).

**Figure 4.**
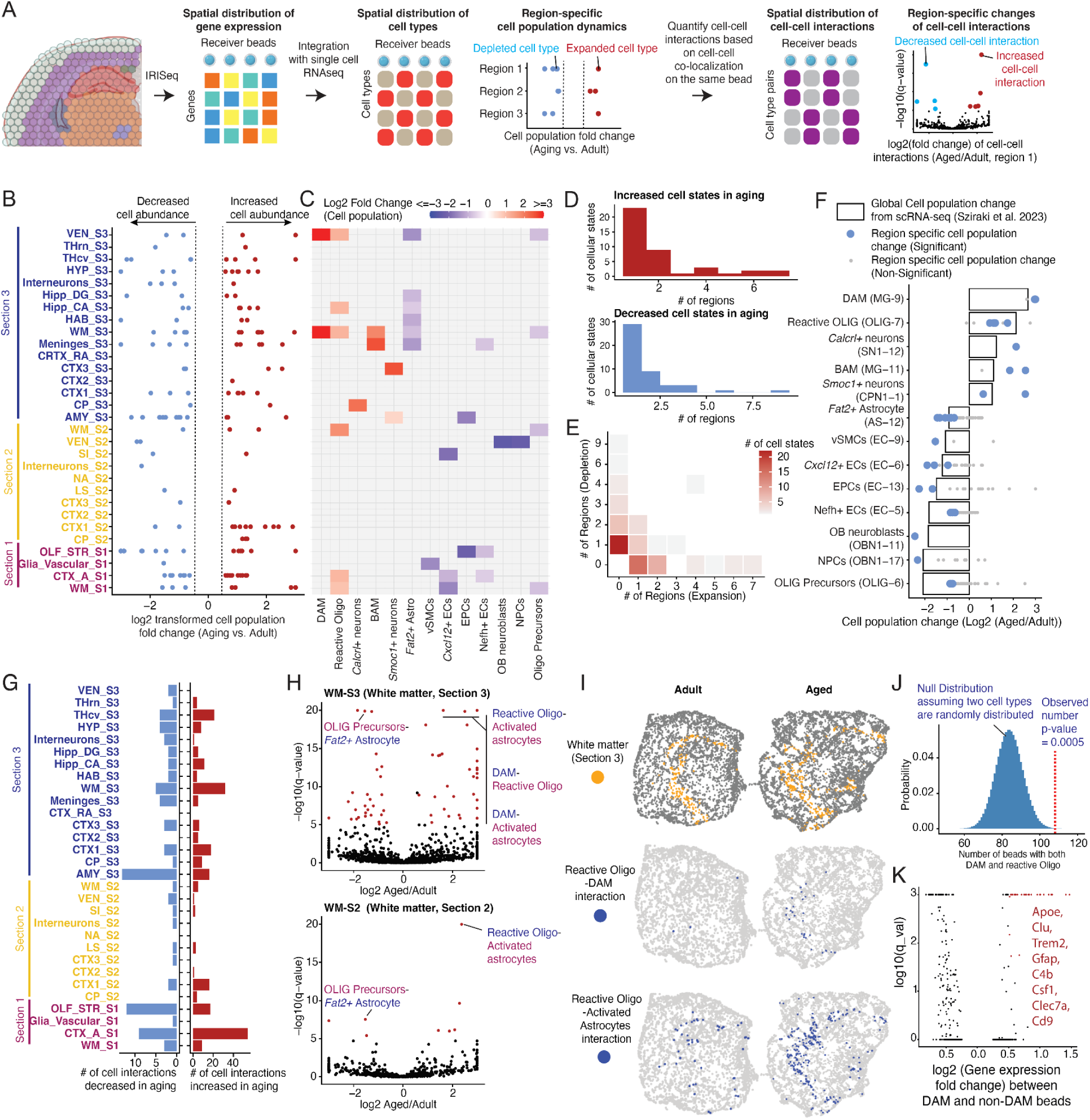
Age-associated changes in region-specific cell type abundance and cell-cell interactions. **(A)** Overview of the approach used to identify region-specific cellular depletion or expansion associated with aging, and to analyze cell-cell interaction dynamics. (**B**) Scatter plot illustrating the number of cell types across different brain regions showing differential abundance between adult and aged animals. Increases are shown in orange and decreases in blue. Differentially abundant cells were defined by a 1.5-fold change with an FDR < 1e-5. (**C**) Heatmap illustrates the fold change in key aging-associated cell types across different brain regions. (**D**) Histograms illustrating the number of cell states that significantly increased (Top) or decreased (Bottom) across different brain regions. The x-axis represents the number of brain regions, while the y-axis shows the count of cellular states. (**E**) Heatmap displays the number of brain regions where cellular states either expand (x-axis) or deplete (y-axis) with aging. The color gradient from light to dark red indicates the number of cell states, with darker shades representing higher numbers. (**F**) This panel illustrates both global (Sziraki et al.) and region-specific changes (IRISeq) in various cell populations with aging. Significant changes are highlighted to show how specific cell types respond to aging in different brain regions, providing insights into the cellular dynamics associated with the aging process. (**G**) Barplot depicting the number of significantly increased and decreased cell-cell interactions associated with aging across different annotated brain regions (Significance FDR of 1e-5 and 1.5-fold change). (**H**) Volcano plot showing significant enrichment and depletion of cell-cell interaction changes in aged versus adult white matter, across two anatomically distinct white matter regions in sections 2 and 3, highlighting significantly changed cell-cell interactions in aging. (**I**) Representative spatial maps depicting white matter regions in two sections from the adult and aged brain. The top panels show representative maps of white matter regions from adult and aged brains. Middle panels display beads indicating colocalization of reactive oligodendrocytes and DAM. The bottom panels show beads with both reactive oligodendrocytes and activated astrocytes. (**J**) Histogram illustrating the null distribution of beads with both cell types assuming random distribution, with a dashed line marking the observed number. (**K**) Volcano plots compare gene expression in beads with or without DAM in the white matter of Section 3, highlighting the top differentially expressed genes.

To analyze age-associated changes in the spatial distribution of cell types, we quantified cell-type-specific fractions across each of the thirty reconstructed brain regions. Using differential abundance analysis between adult and aged conditions (**Methods**), we identified 170 region-specific differentially abundant cellular subtypes (1.5-fold change between ages with an FDR of 1e-5) (Figure 4B-C, **Table S6**). Notably, these aging-associated subtypes are largely altered in one or two unique regions (Figure 4D). For cell types that changed in multiple regions, their expansion or depletion was consistent across regions (Figure 4E). Moreover, despite the deconvolution pipeline not accounting for aging-associated cell dynamics, a comparison between the top region-specific cell population changes in the *IRISeq* spatial dataset and the published scRNA-seq dataset (*20*) showed high consistency, validating the reliability of our cell type deconvolution pipeline and regional cell abundance analysis (Figure 4F).

Compared to single-cell analysis, spatial transcriptome analysis offers unique advantages for identifying region-specific cell population alterations in the aged brain. For example, the most depleted populations in the aged brain included OB neuroblasts (OBN 1-11, marked by *Prokr2* and *Robo2*(*38, 39*)) and OB neuronal progenitor cells (OBN 1-17, marked by *Mki67* and *Egfr*(*40*)), both detected primarily in the reconstructed ventricle regions from section 2 (**Fig. 4C**). This aligns with the well-known location of adult neurogenesis along the walls of the lateral ventricles (*28*). In contrast, oligodendrocyte precursor cells (OLG-6, marked by *Prom1* and *Tcf7l1* (*40*, *41*)) were depleted in aged white matter and ventricle regions, indicating impaired oligodendrocyte differentiation with age (**Fig. 4C**). Additionally, various vascular and blood cell types, including *Myocd+* vascular smooth muscle cells (*Myocd+* vSMC, EC-9), *Cxcl12+* endothelial cells (EC-6), *Nefh+* endothelial cells (EC-5), and *Hbb-bs+* erythrocyte progenitor cells (EC-13) were significantly depleted (**Fig. 4C**), consistent with age-related declines in blood flow, energy supply, and neurovascular functions (*42*). The depletion of these cell types corresponds with global cell population changes (**Fig. 4F**), indicating region-specific dysregulation of the vascular system in aging.

Meanwhile, several cell subtypes significantly expanded across different regions in the aged brain (**Fig. 4B**-**C**). These included a microglia subtype (MG-9, *Apoe+, Csf1+*), identified as disease-associated microglia (’DAM’)(*43*), and a reactive oligodendrocyte subtype (OLG-7, *C4b+, Serpina3n+* (*44*, *45*)), both predominantly found in the white matter and ventricle regions, validated by the *10x Visium* dataset (*20*) (**Fig. 4C**, **Fig. S8A-B**). This expansion aligns with the notably upregulated inflammatory response in these areas (**Fig. 3D**). Furthermore, we observed a region-specific expansion of *Lyz2+* border-associated macrophages (*Lyz2+* BAM, MG-11) around the meningeal regions (**Fig. 4C**). This cell type is known to degrade the extracellular matrix, contributing to vascular function defects (*46*, *47*), which is consistent with the observed depletion of endothelial cells in the same region (**Fig. 4C**).

Leveraging the *IRISeq* platform, where each bead captures transcriptomes from multiple cells within the same neighborhood, we quantified the colocalization of two cell subtypes around the same beads to assess local cellular spatial interactions (**Methods**). We next conducted a differential analysis to identify cell-cell spatial interactions that are significantly altered with aging across various brain regions. This analysis revealed a total of 499 cell-cell spatial interaction pairs that are either significantly upregulated or downregulated in aging (1.5-fold change between ages with an FDR of 1e-5), prominently in regions such as the cortical associated area, olfactory/striatal area, amygdala, and white matter region (**Fig. 4G**, **Table S7**).

While most aging-associated changes in cell-cell interactions are highly region-specific (**Fig. S9A**), we identified several interaction pairs that consistently occur across multiple regions (**Fig. S9B**). For example, in the aged white matter, we observed a significant decrease in interactions between oligodendrocyte precursor cells (OLG-6) and *Gfap*-low *Fact2*-high astrocytes (Astrocyte-12) across different sections (**Fig. 4H**, **Fig. S9C**), corresponding with their reduced abundance in these regions during aging (**Fig. 4C**). Conversely, we noted a notable increase in interactions among three cell types: disease-associated microglia (‘DAM’, MG-9), reactive oligodendrocytes (OLIG-7), and *Gfap*-high activated astrocytes (Astrocyte-9), which are involved in astroglial activation and gliosis during aging and neurodegeneration (*48*) (**Fig. 4H**-**I**, **Fig. S9C**). The up-regulated co-localization of these cell types in the aged white matter could form a potential “DAM niche”, which we further validated by the *10x Visium* technique (**Fig. S9D**).

To assess whether the increased spatial interaction of these cell subtypes is simply due to their expansion, we performed a hypergeometric test comparing the observed number of beads with colocalized cell types against a null model assuming random distribution within the region. The results demonstrate that the interactions between reactive oligodendrocytes, DAM, and activated astrocytes are not only more frequent in the aged brain but are also significantly enriched within the same neighborhood (p-values = 0.0005 and 1e-17 for interactions between DAM and reactive oligodendrocytes, and between DAM and activated astrocytes) (**Fig. 4J**, **Fig. S9E**). As a further validation, gene signature analysis of beads within the “DAM niche” shows significantly higher expression of markers for activated astrocytes (*e.g., Gfap*) and reactive oligodendrocytes (*e.g., C4b*) compared to non-DAM beads from the same region (**Fig. 4K**), underscoring altered cellular interaction and communication networks as key features of brain aging.

### Discussion

The rapid advancement of spatial transcriptomics technologies has revolutionized our ability to map the locations and interactions of heterogeneous cellular states in situ(*1–10*), particularly in the context of aging or disease. However, most current approaches, especially imaging-based methods, often prioritize resolution at the expense of throughput and time, typically restricting analyses to specific tissue regions and limiting their utility across various conditions and replicates.

To address these limitations, we introduce *IRISeq*, an optics-free spatial profiling platform that leverages “spatial interaction mapping by indexed sequencing”. *IRISeq* bypasses the need for imaging or pre-indexed arrays, allowing multiple sections to be processed in a single day at significantly reduced costs—approximately $30 per tissue section—without requiring complex equipment. Furthermore, its optics-free design is particularly suited for mapping large tissue areas exceeding 1cm², which are traditionally limited by imaging speed and the constraints of pre-indexed arrays. Additionally, *IRISeq* offers adjustable resolution based on bead size (ranging from 5 to 50 µm), enhancing its flexibility and user-friendliness. These features make *IRISeq* a versatile and accessible tool for a wide range of applications, from basic research to clinical diagnostics.

Using the *IRISeq* platform, we explored age-related changes in gene expression and cellular dynamics across thirty regions of the mouse brain. Our findings indicate a general down-regulation of genes associated with mitochondrial activity, ribosomal function, and neuronal modulation across almost all regions. Conversely, genes related to the complement and interferon pathways, as well as inflammation markers, were predominantly up-regulated in specific areas such as the ventricles and white matter. This regional specificity aligns with previous reports highlighting increased susceptibility to aging in these locations(*34*, *49*).

Further cell-centric analysis pinpointed precise region-specific changes in cell populations. For example, we observed a marked depletion of neurogenesis-related cells in the subventricular zone and oligodendrocyte precursor cells in the white matter and ventricles, corroborating earlier studies(*50*). Additionally, our analysis uncovered significant changes in other cell types, such as the expansion of border-associated macrophages (BAM) and the concurrent depletion of *Gfap*-low *Fact2*-high astrocytes and vascular endothelial cells in the aged meninges. Moreover, we developed a computational pipeline to analyze local cell-cell interactions by assessing the co-localization of two cell types on the same beads. This approach revealed that in the aged white matter, not only is there an expansion of disease-associated microglia, reactive oligodendrocytes, and activated astrocytes, but there is also a significant increase in their co-localization, independent of changes in their abundance. This observation aligns with findings from previous imaging-based studies(*34*) and validates the effectiveness of our platform in detecting region-specific alterations in cellular networks during aging.

*IRISeq* is a highly optimized platform for spatial transcriptomics, yet there are opportunities for further improvement. Currently, the method does not provide isolated single-cell information. However, it is readily compatible with nuclei hashing techniques that can extend its capabilities to direct spatial analysis at the single-cell level (*51*). Additionally, *IRISeq* can be integrated with methods to profile proteins(*52*) and epigenetic landscape(*53*), broadening its application spectrum. A current limitation lies in the image reconstruction process, which relies on the stochastic UMAP algorithm(*19*). This results in digital approximations that do not exactly replicate the original images. Despite this, the technique robustly maps the relative locations of different anatomical regions, as confirmed through comparisons with the *10x Visium* platform. It’s important to note that our downstream biological analyses—including transcriptome annotation, cell population dynamics, and cellular interaction studies—remain unaffected by these imaging constraints.

In summary, *IRISeq* demonstrates significant potential for detailed mapping of region-specific molecular signatures, cell population dynamics, and local cell-cell interactions across varied and complex biological landscapes. Its capacity to reconstruct images based solely on sequencing local DNA interactions allows for profiling of tissues without size constraints and across varied resolutions. Looking ahead, the high-throughput, cost-effective nature of *IRISeq* positions it as a transformative tool for comprehensive spatial mapping of entire organs or organisms, across various genders, ages, and disease states. This opens new possibilities for identifying region-specific vulnerabilities linked to different diseases, enhancing our understanding of complex biological systems.

## Acknowledgments

We thank members of the Cao lab and Center for Integrated Cellular Analysis (CEGS) for helpful discussions and feedback. We also thank members from the Rockefeller University facilities, specifically the Comparative Bioscience Center and High-Performance Computing Center for their support with animal maintenance and data analysis.

## Funding

This work was funded by grants from the NIH (1DP2HG012522, 1R01AG076932, and RM1HG011014 to J.C). W.Z. was funded by the Kellen Women’s Entrepreneurship Fund. A.A. was supported by a Medical Scientist Training Program grant from the National Institute of General Medical Sciences of the National Institutes of Health under award number: T32GM152349 to the Weill Cornell/Rockefeller/Sloan Kettering Tri-Institutional MD-PhD Program.

## Author contributions

J.C. and W.Z. conceptualized and supervised the project. A.A. designed molecular biology and library preparation experimental steps with help from W.J. A.A. and W.J. performed optimizations for the experimental pipeline with input from Z.Z, Z.X. and T.R. W.J. processed the mice brain samples for the *IRISeq* profiling experiment, with the help of A.A. A.A. performed computational analyses with input from J.C. and W.Z. S.I. and A.A. performed website development and online visualization. W.Z., J.C., and A.A wrote the manuscript with input and biological insight from all co-authors.

## Data and materials availability

A detailed protocol of the *IRISeq* platform is outlined in **supplementary note 1**. Raw FASTQ files, processed count matrices, cell metadata, and gene metadata can be downloaded from NCBI GEO under accession number GSE270383 (GEO reviewer token: **izwrekiyffwjhyp**)

## Materials and Methods

### Animals

C57BL/6 wild-type mice were acquired from the Jackson Laboratory and the National Institute on Aging colony at Charles River. All mice were housed under standard conditions, with groups matched for sex and age. Brain tissues were collected from mice at 23 months (n=3) and 4 months (n=3), including three females in each group. Mice were socially housed. All animal procedures complied with institutional, state, and federal regulations, and were approved under IACUC protocols 21049 and 20047. Comprehensive metadata for each animal, including individual ID, sex, age, birth, and euthanasia dates, are detailed in **Table S2**.

### IRISeq library preparation

Detailed step-by-step *IRISeq* protocol is included as a supplementary protocol (**Supplementary Note 1**). **Computational procedures for processing *IRISeq* libraries**

### Beads Connections Processing

Following the sequencing process, data from two beads—receiver beads and sending beads—were obtained, with unique molecular identifiers (UMIs) used to identify and quantify individual connection molecules. Reads were classified into R1 (receiver bead barcodes) and R2 (sender bead barcodes), and mapped to a whitelist of barcodes with a 1 base pair error tolerance. Duplicated reads were filtered out based on UMIs, and a CSV file was generated containing the receiver bead barcode, UMI, and sender bead barcode, where each line represented a unique molecular connection between beads.

To ensure data quality, connections between beads with fewer than 7 UMIs were filtered out. A matrix was then generated, with each row representing a receiver bead, each column representing a sender bead, and the values corresponding to the number of UMIs for each connection. Principal Component Analysis (PCA) was applied to this matrix to reduce dimensionality while preserving variance, using an elbow plot to determine the optimal number of principal components, typically capturing around 80% of the variance. Subsequently, Uniform Manifold Approximation and Projection (UMAP) was performed on the PCA matrix to further reduce dimensionality, using parameters such as a minimum distance of ∼0.2 and approximately ∼20 neighbors to obtain a square-shaped projection. UMAP coordinates for each receiver bead were saved in CSV files for spatial mapping. For larger arrays, UMAP was run on a GPU with increased training epochs (from ∼500 to ∼10,000 or more) to handle the larger data matrix. Additionally, for large array analysis, density-based spatial clustering of applications with noise (DBSCAN) was applied post-UMAP to remove several erroneously mapped beads (*54*). A detailed workflow is summarized in **Figure S2**.

### cDNA Data Matrix processing

After the sequencing process, the data is obtained as two sets of fastq files. The first set of fastq files (R1.fastq) contains the barcode sequences for receiver beads along with their associated UMIs, while the second set of fastq files (R2.fastq) contains the tagmented cDNA sequences. The barcode sequences from R1.fastq are mapped to a whitelist of bead barcodes with a tolerance of one base pair error, and the identified bead barcode is added as an identifier for the corresponding sequences in R2.fastq. New Read2 files are then generated, incorporating the corresponding Read1 barcodes in the sequence names, and any PolyA tails present at the end of the sequences are trimmed for accurate mapping and analysis. The filtered and trimmed reads are mapped to the reference genome using STAR (Spliced Transcripts Alignment to a Reference) (*55*) to align the sequences from R2 to the genomic sequences, identifying their locations and potential splice sites. Duplicate reads are identified and removed post-mapping to eliminate redundant data and ensure accuracy in downstream analysis, resulting in new SAM files containing the mapped reads with duplicates removed. Finally, gene count matrices are generated from the processed SAM files for each receiver bead, quantifying the number of reads aligned to each gene and providing information about gene expression levels.

### Dimensionality Reduction, Clustering, and Region Annotation

Raw data from the cDNA sequencing experiment, presented as receiver bead-by-gene expression matrices, undergoes several preprocessing steps. The first step includes removing Beads with low UMI counts (Less than 600 for the mouse aging dataset). The second step includes dividing the data set according to the anatomical sections by which tissue sections were cryosectioned. *LIGER* objects are constructed by first normalizing the number of UMIs (*23*). Then 2000 highest variable features are selected for each tissue section, followed by scaling of gene expression features. After data normalization and processing, each anatomical data set with associated individual tissue sections is integrated by *LIGER*’s non-negative matrix factorization (iNMF) approach, followed by UMAP mapping. Louvain clustering is then performed on the normalized NMF factor loadings. Clusters with low UMI and non-specific gene features were iteratively removed to ensure data quality, followed by *LIGER* iNMF workflow to recluster and run UMAP again until no low-quality clusters were formed.

Brain region identification was performed based on NMF factors gene markers, and also performing Wilcoxon differential expression analysis for each identified cluster. A similar analysis scheme and data processing were performed for *IRISeq* and *10X Visium* integration analysis, and *10X Visium* aging validation analysis. Similar normalization and scaling were done for *IRISeq* high-resolution data analysis.

Large-area dataset analysis (see in Figure 1) was performed utilizing Scanpy(*56*) for principal component analysis (PCA) followed by UMAP. Leiden clustering was implemented using Scanpy(*56*) to group beads with similar gene expression profiles. Additionally, beads within each cluster are annotated by utilizing differential gene expression analysis for each cluster, and associated cluster-specific genes to known cell type and/or brain regions, along with utilizing genes and/or cluster mapping onto spatial anatomical location.

### Differential Expression Analysis

In our differential expression (DE) gene analysis, we employed the likelihood ratio test to identify aging differential expression for specific regions using Monocle2(*57*). DE genes were filtered based on the following cutoffs: q-value < 0.05, with fold change (FC) > 3 between the maximum and second expressed condition, and with transcripts per million (TPM) > 50 in the highest expressed condition.

### Cell Type Deconvolution

To perform cell type deconvolution, we applied robust cell type decomposition (*RCTD*) (*25*), a computational method that uses cell type profiles from single-cell RNA-seq to decompose cell-type mixtures while accounting for differences across sequencing technologies. We integrated *IRISeq* spatial expression data with a single-cell transcriptome dataset from an earlier study (*20*) following parameters suited for ∼50 µm-sized spots and excluded cell types not present in the microdissected sections.

### Differential Abundance Analysis

To perform cell abundance analysis across different regions, first, *RTCD* maximum-likelihood cell type proportions are binarized by assigning them into one of five categories: values less than 0.05 were categorized as 0, values between 0.05 and 0.2 as 1, values between 0.2 and 0.6 as 2, values between 0.6 and 0.8 as 3, and values greater than 0.8 as 4. This method allowed for the transformation of continuous proportion data into discrete categories, facilitating subsequent analysis and visualization. After this, a count matrix of receiver bead by cell subtypes is obtained. We later counted the number of cells for each cell subtype and normalized these counts against the total cell number. For differential abundance analysis of cell subtypes, we used the negative binomial model, a common approach for differential gene expression analysis that is well-suited for count data and large-scale datasets. We used a likelihood-ratio test to identify differentially abundant cell subtypes, employing the differentialGeneTest() function of Monocle2 (version 2.28.0) (*57*).

For fold change calculations, we first normalized the cell counts for each cell subtype relative to the total cell count in each condition. We then compared these normalized values between case and control conditions, adding a small numerical value (10^−6) to reduce noise from very small clusters. To classify a cell subtype as a “significantly changed cell subtype,” we set criteria of a maximum false discovery rate of 1e-5 and a fold change greater than 1.5 between conditions. We quantified the abundance of aging-associated cell subtypes in the different annotated brain regions by performing differential abundance tests comparing adult and aged samples. Recognizing that variations in one cell subtype can influence the relative proportions of others, particularly static cell subtypes, our analysis focused on cell subtypes exhibiting significant changes—more than a 1.5 shift in population during aging. This approach is based on the observation that many significantly changed cell clusters correspond to rare cell states and represent a small portion of the global cell population. Consequently, even if these changing cell subtypes have a substantial impact on others, we expect the overall relative proportion shifts to remain within a two-fold range.

### Cell-cell interaction analysis

To assess whether the increased spatial interaction of specific cell subtypes is due to their expansion, we performed a hypergeometric test. This test compared the observed number of beads with colocalized cell types against a null model that assumes a random distribution of cell subtypes colocalization within the region. Prior to conducting the hypergeometric test, we created a new receiver bead by cell-cell interaction matrix. We generated this matrix by first filtering out low RTCD likelihood probability values (< 0.05) for subtype deconvolutions. Next, we conducted a pairwise analysis of cell subtype co-existence on the same bead. If two cell subtypes had a positive value after filtering out low RTCD likelihood probability values, we added a count of 1 to the corresponding cell-cell interaction column. Using the hypergeometric test we defined the total number of beads within the region (N), the number of beads with a specific cell subtype-subtype interaction (k), the number of beads where colocalization is observed (n), and the number of beads with the other interacting cell subtype (x). The hypergeometric probability *P* (*X* = *x*) was calculated as:

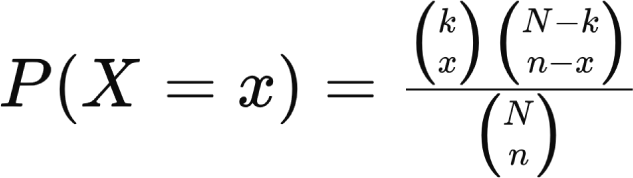

This formula determined the likelihood of observing the number of colocalized cell subtypes under the null hypothesis.

## Supplementary Text

**Fig. S1.**
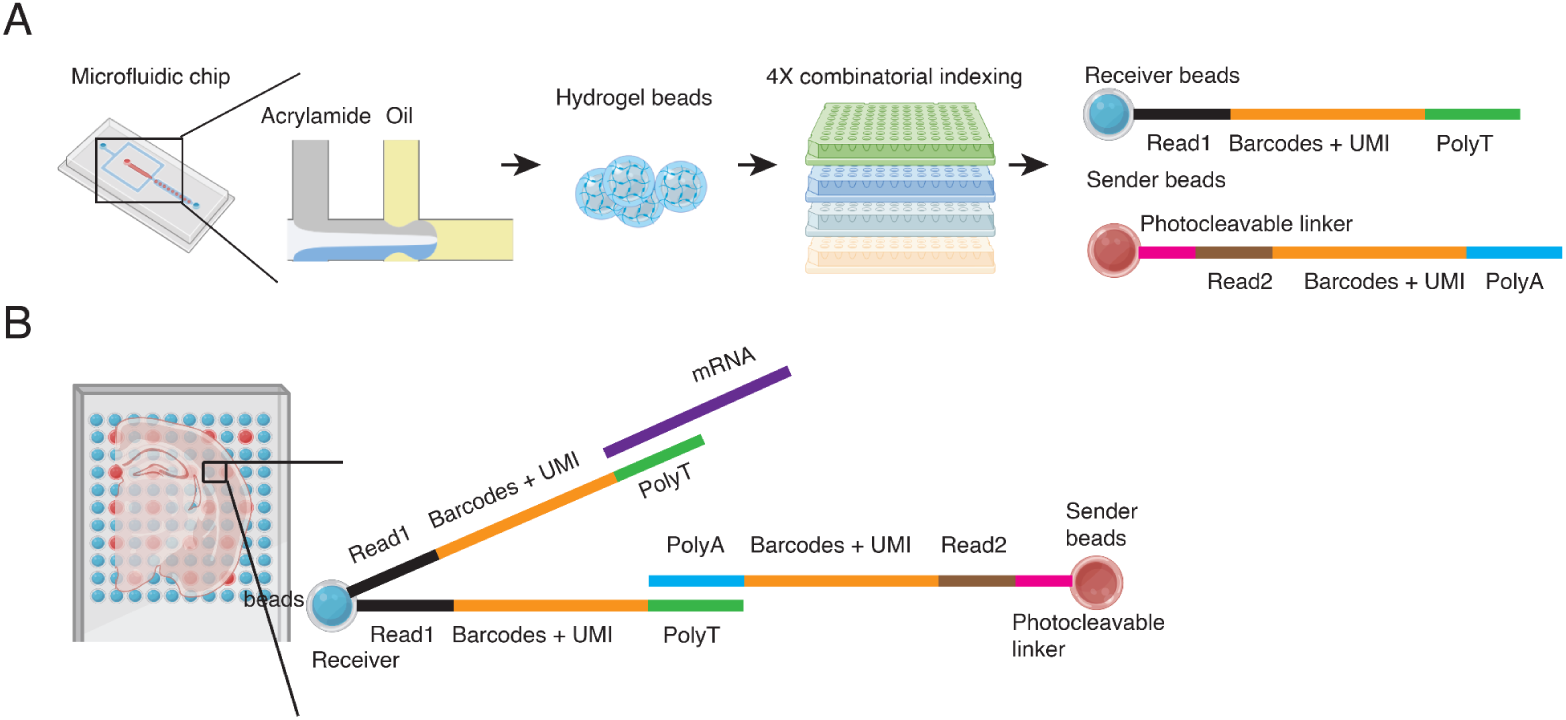
Bead Generation and Barcoding in *IRISeq*. (**A**) Schematic illustrating the generation of barcoded hydrogel beads (‘sender beads’ and ‘receiver beads’) through combinatorial indexing. (**B**) Diagram showing the dual interaction between sender and receiver beads for capturing RNA from tissue samples and indexing spatial locations on the array.

**Fig. S2.**
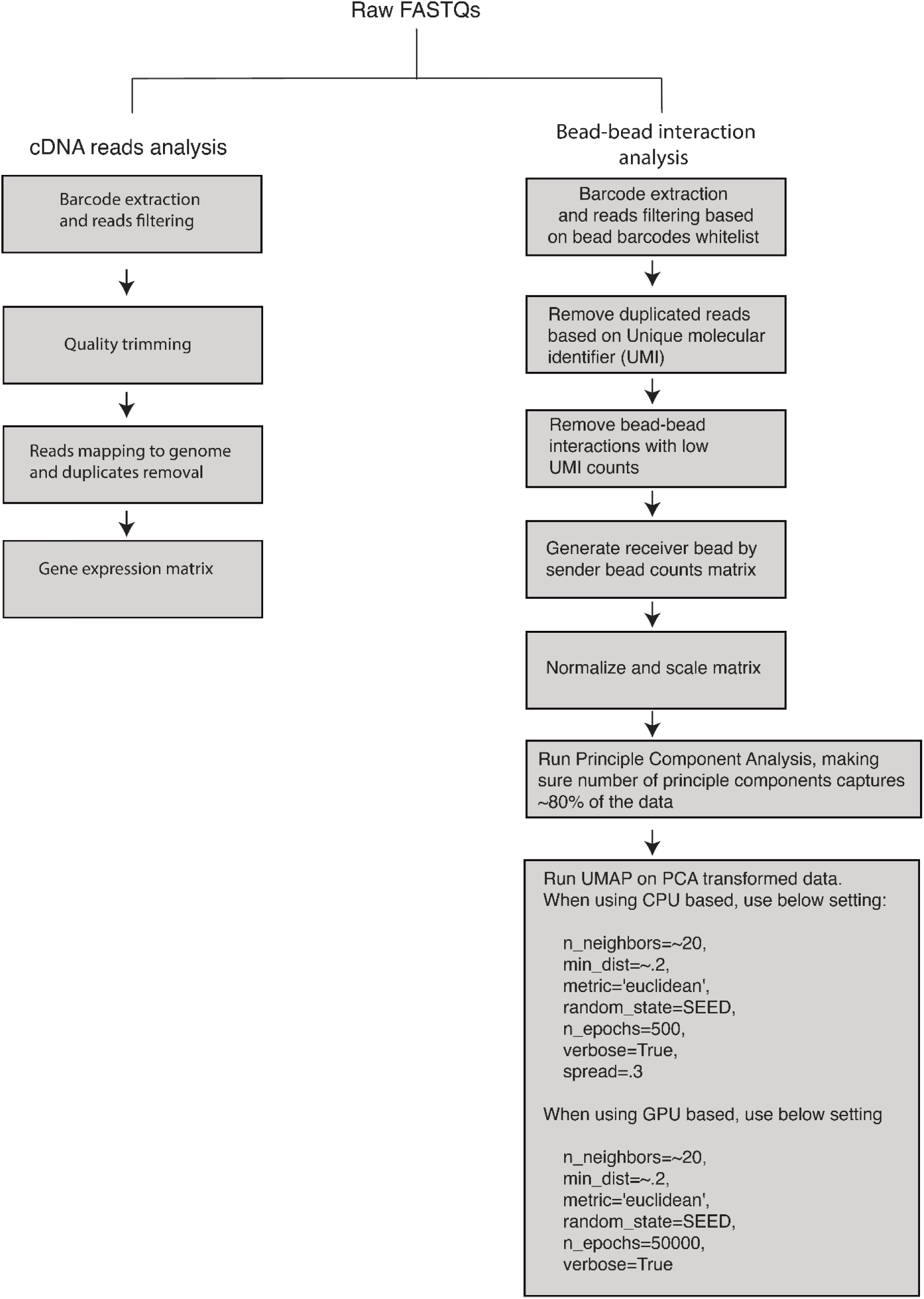
*IRISeq* computational pipeline. Customized computational pipeline outlining the derivation of two matrices from each *IRISeq* experiment. The gene count matrix captures gene expression for each bead, while the bead-bead connection counts matrix infers bead locations for image reconstruction.

**Fig. S3.**
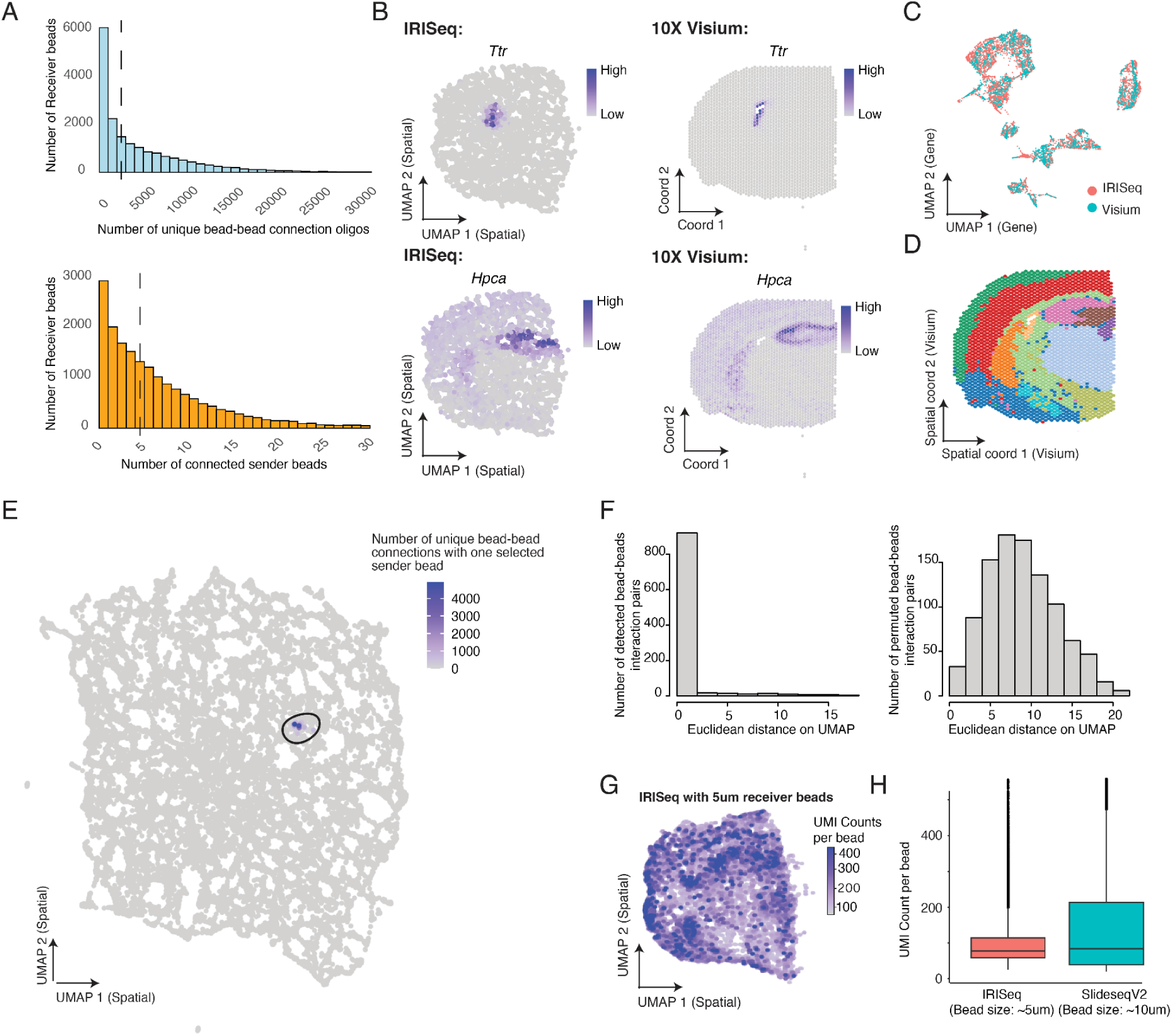
*IRISeq* quality control and performance comparison. (**A**) The top panel displays the distribution of unique bead-bead interactions per receiver bead. The bottom panel shows the distribution of connected sender beads per receiver bead, with the median indicated by a dashed line in both panels. (**B**) UMAP plots compare the spatial distribution of receiver beads from a small-scale *IRISeq* experiment (left) with *10x Visium* data (right), colored by the expression of region-specific markers *Ttr* (ventricle region, top) and *Hpca* (hippocampus region, bottom). **(C**) UMAP plot showing the integrated gene expression profiles from *IRISeq* receiver beads and *10x Visium* spots, colored by assay type. (**D**) UMAP plot illustrating spatial distribution colored by annotated brain regions, similar to Figure 1G. (**E**) UMAP plot showing the reconstructed spatial distribution of receiver beads from *IRISeq*, with beads colored according to the number of unique interaction oligos with a selected sender bead, validating the spatial reconstruction pipeline. (**F**) Histograms depicting *Euclidean* distances (in Figure S3E) between pairs of receiver beads connected to the same sender bead (left) versus distances between randomly selected receiver beads (right). (**G**) UMAP plot visualizing the spatial distribution of 5 µm receiver beads in a high-resolution *IRISeq* experiment, colored by the number of unique transcripts per bead. (**H**) Boxplot comparing the number of unique transcripts per bead in *IRISeq* (based on 5 µm beads) versus *Slide-seqV2* data (based on 10 µm beads).

**Fig. S4.**
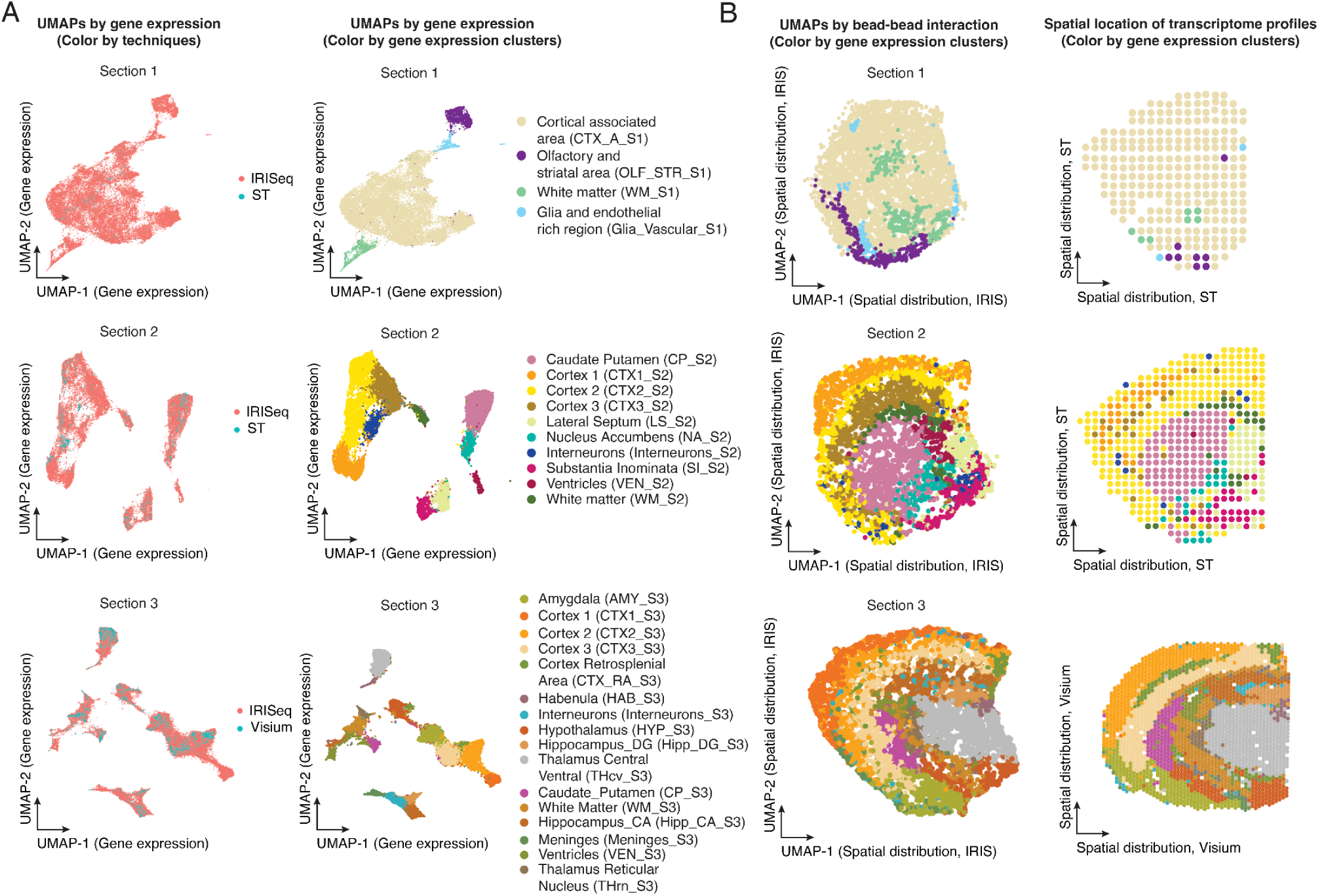
*IRISeq* region annotation and published datasets comparison. (**A**) UMAPs by gene expression comparing *IRISeq* data with published spatial transcriptomics datasets (*24*). The left panels show the gene expression profiles colored by techniques (*IRISeq*, ‘*ST’ for spatial transcriptomics, ‘Visium’ for 10X Visium*), while the right panels color the beads by gene expression clusters specific to each brain region. (**B**) The left panels depict UMAPs showing the reconstructed spatial distribution of receiver beads from *IRISeq*, colored by gene expression clusters. The right panels display the spatial distribution of transcriptome profiles from published datasets (*20*, *24*) (‘*ST’ for spatial transcriptomics, ‘Visium’ for 10X Visium*), with coloring corresponding to the annotated brain regions as identified in the gene expression clustering in (A).

**Fig. S5.**
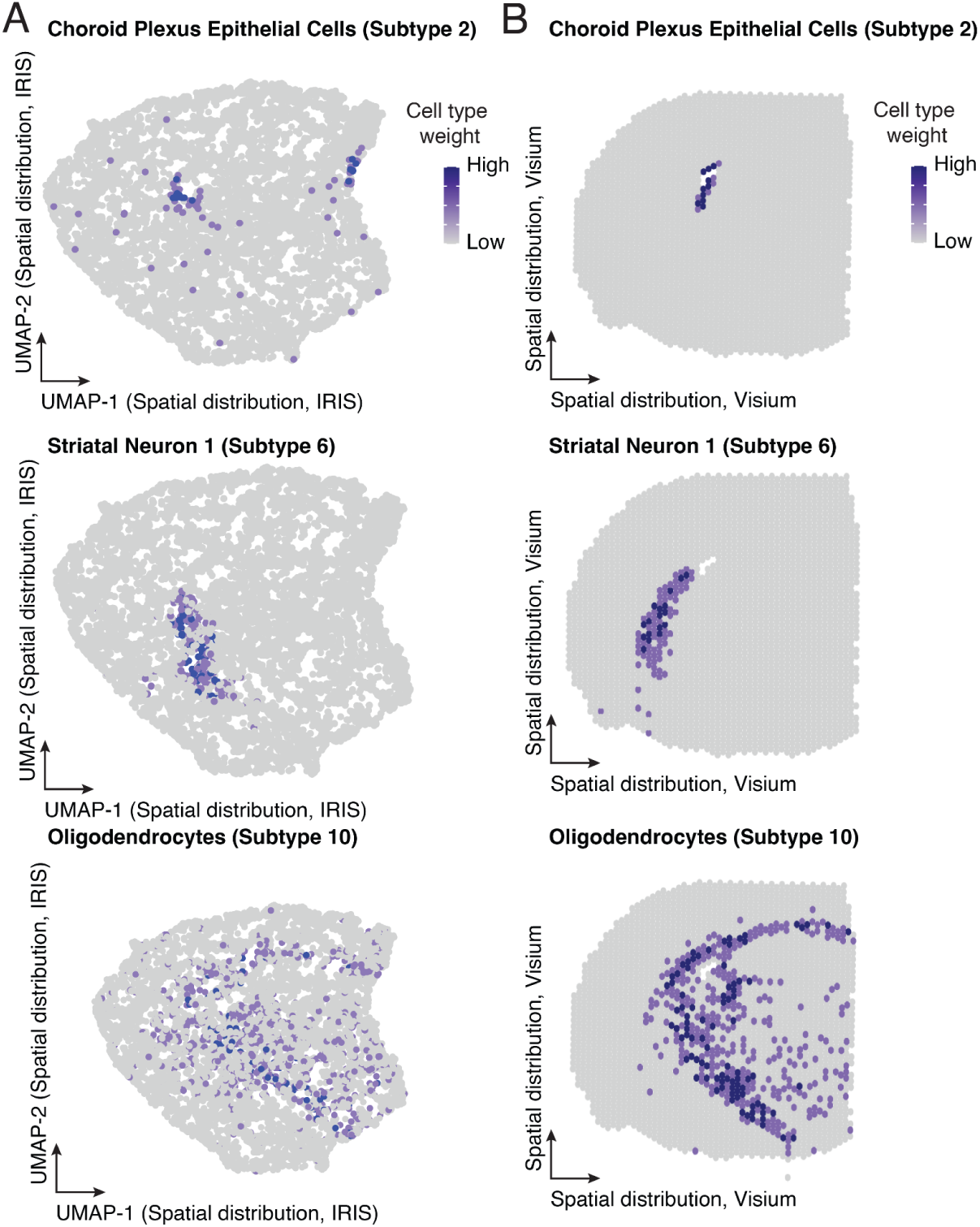
Comparison of cell subtype distributions between *IRISeq* and *10X Visium*. (**A**) UMAP plots showing the reconstructed spatial distribution of receiver beads from *IRISeq*, colored by cell type weight. Each plot highlights specific cell types, including Choroid Plexus Epithelial Cells (Subtype 2), Striatal Neuron 1 (Subtype 6), and Oligodendrocytes (Subtype 10), indicating the abundance of these cell types as determined by RTCD analysis(*25*). (**B**) Corresponding UMAP plots showing the spatial distribution of transcriptomes generated using *10X Visium* (*20*), also colored by cell type weight for the same cell types as in panel (A).

**Fig. S6:**
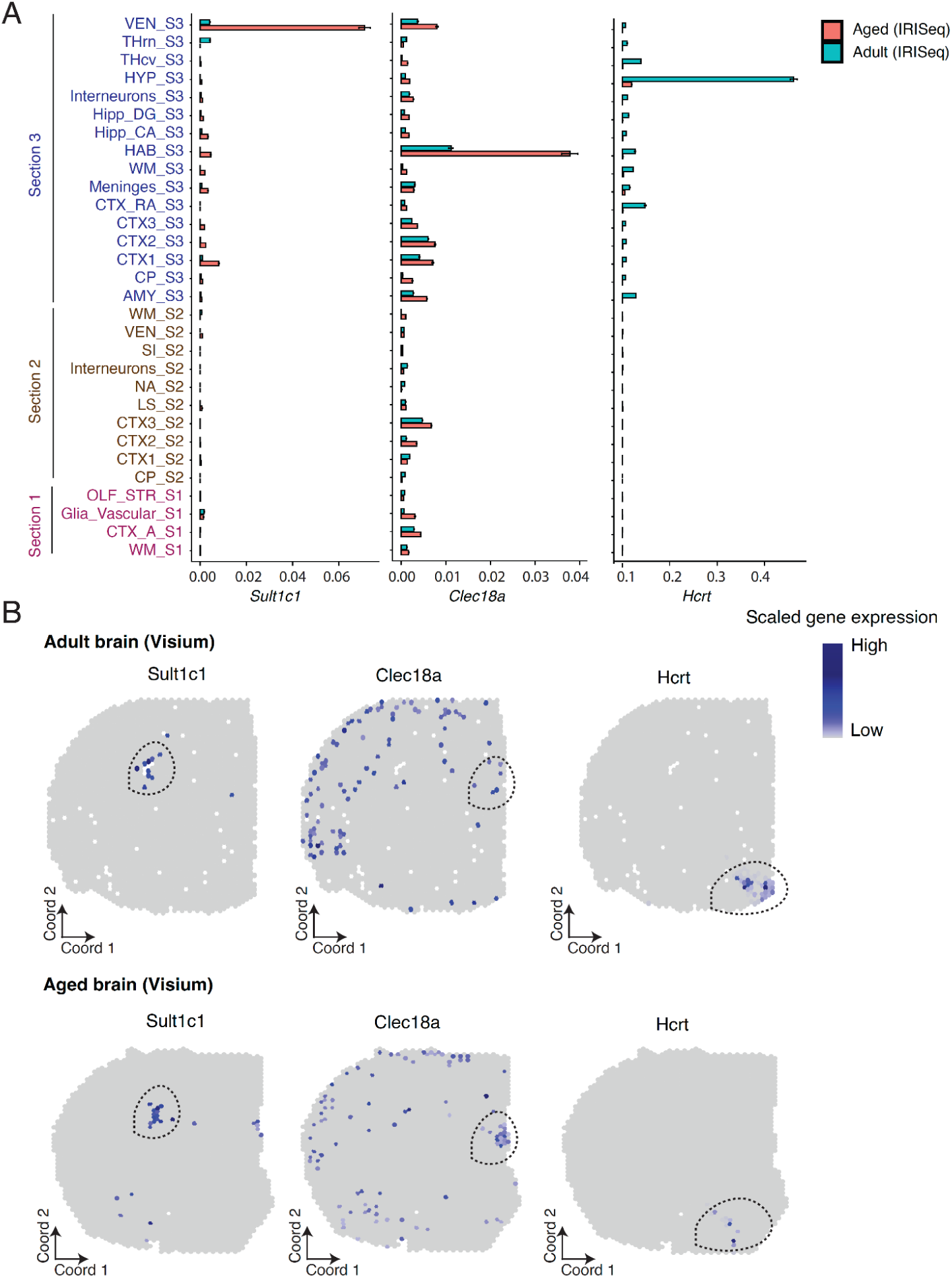
Region-specific gene expression changes in aging. (**A**) Bar plots displaying the scaled expression and standard error of selected genes (*Sult1c1, Clec18a, Hcrt*) across brain regions, comparing adult (blue) and aged (red) conditions. The expression values are normalized by the total UMI count per bead and log-transformed. (**B**) Spatial distribution of transcriptomes from published *10x Visium* data(*20*), depicting gene expression in adult (top) and aged (bottom) brain sections. Maps are colored by the normalized expression of genes selected in (A), highlighting areas of interest where these genes are predominantly expressed.

**Fig. S7.**
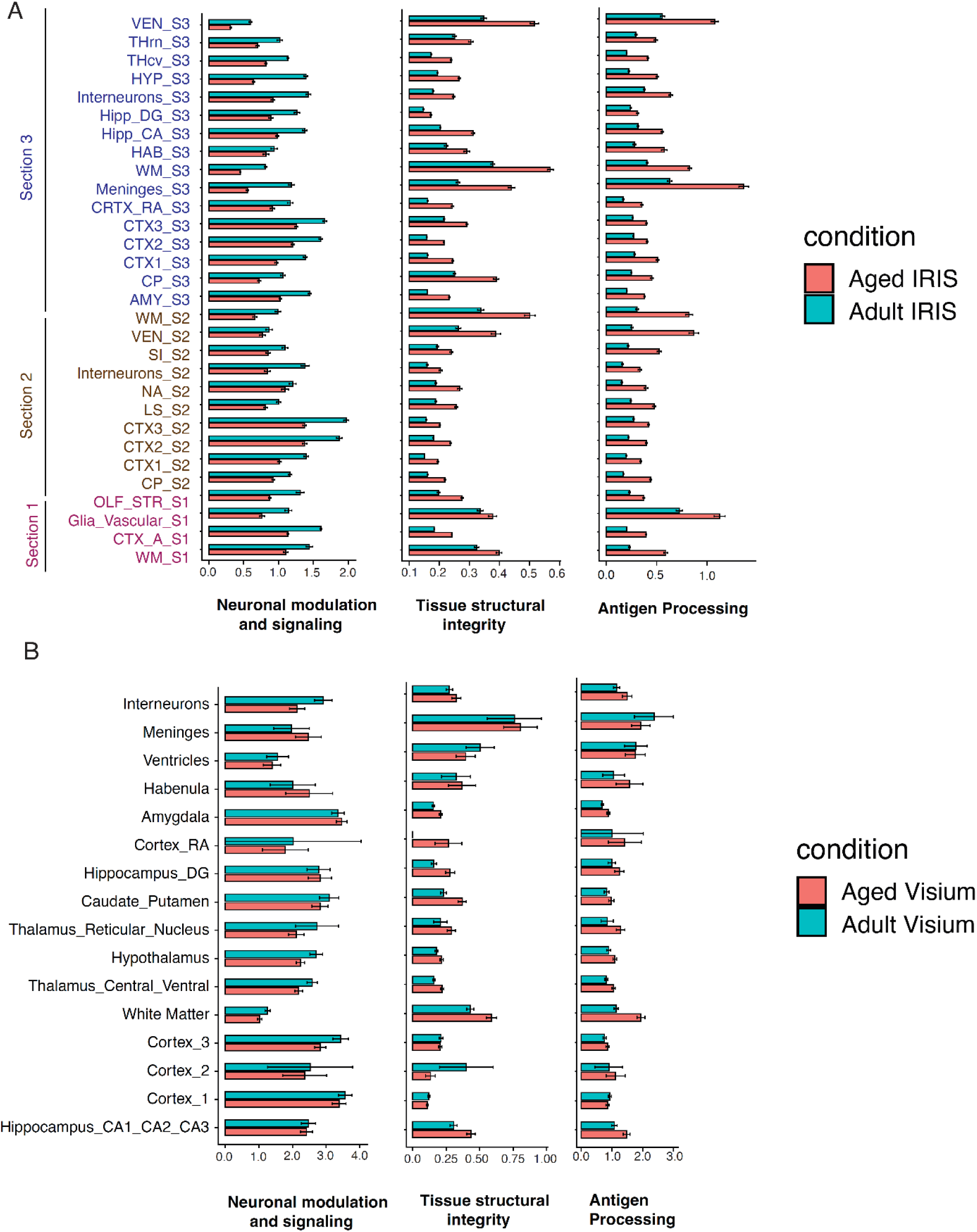
Downregulated and upregulated gene modules exhibiting consistent aging-associated changes across multiple brain regions (A) Bar plots display the scaled expression and standard error for gene modules associated with the neuronal modulation and signaling, tissue structural integrity, and antigen processing pathways across various brain regions in adult (blue) and aged (red) conditions. Specific genes within each module include Neuronal modulation and signaling -[*Sez6, Nts, Asic2, Pdyn, Npy, Cck, Rab37, Lrrc7, Slc32a1, Tac2, Gng13, Npy, Syt17, Hcrt],* Tissue structural integrity *-[Col6a1, Col6a2, Col9a2, Vim, Hapln2*], and Antigen Processing -[*Psmb8, Psmb9, Ctss, H2-K1, H2-Q7, H2-D1, H2-Eb1, B2m, Cd74, Tap1, Tap2*]. (B) Similar bar plots show expression levels derived from published 10x Visium data (*20*), comparing adult (blue) and aged (red) samples across corresponding brain regions. For both (A) and (B), expression values were aggregated from pathway-related genes, normalized, log-transformed, and scaled to facilitate direct comparison across conditions and regions.

**Fig. S8.**
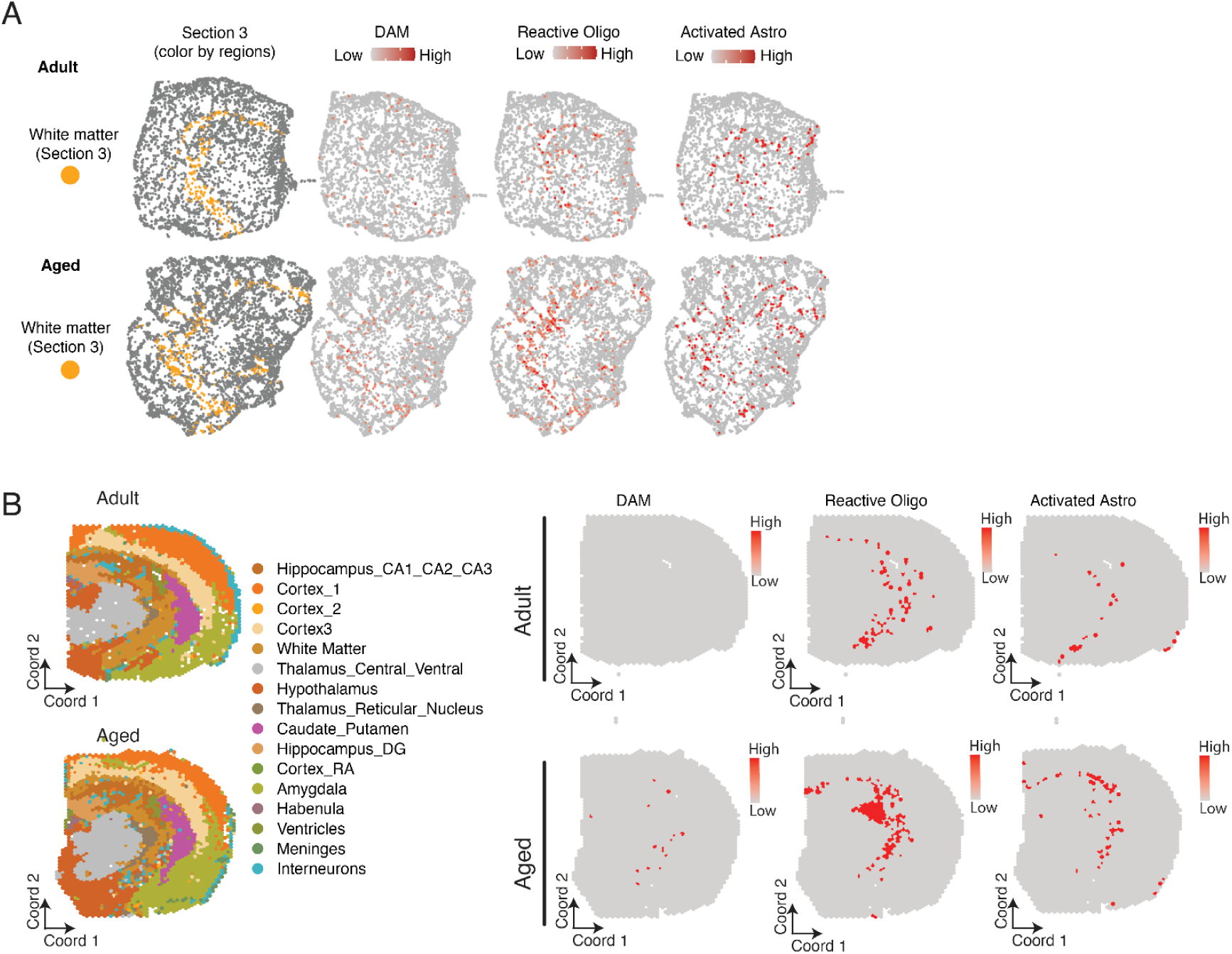
Region-specific alterations of cellular abundance in brain aging. (**A**) Representative spatial maps depicting white matter regions in Section 3 of both adult and aged brains (Left). The right panels display beads colored by cell type-specific weight from *RTCD* analysis, which integrates *IRISeq data* with a single-cell transcriptome atlas. From left to right, the maps show overall white matter localization and distributions of disease-associated microglia (‘DAM’), reactive oligodendrocytes (‘Reactive Oligo’), and activated astrocytes (‘Activated Astro’) in adult (top row) and aged (bottom row) conditions. (**B**) The left panel illustrates the spatial distribution of transcriptomes across various brain regions in the adult and aged brain, color-coded by region. The right series of panels display the distribution of DAM, reactive oligodendrocytes, and activated astrocytes, colored by their relative abundance from *RTCD* analysis that integrates *Visium* data (*20*) with the single-cell transcriptome atlas, shown for both adult and aged samples.

**Fig. S9.**
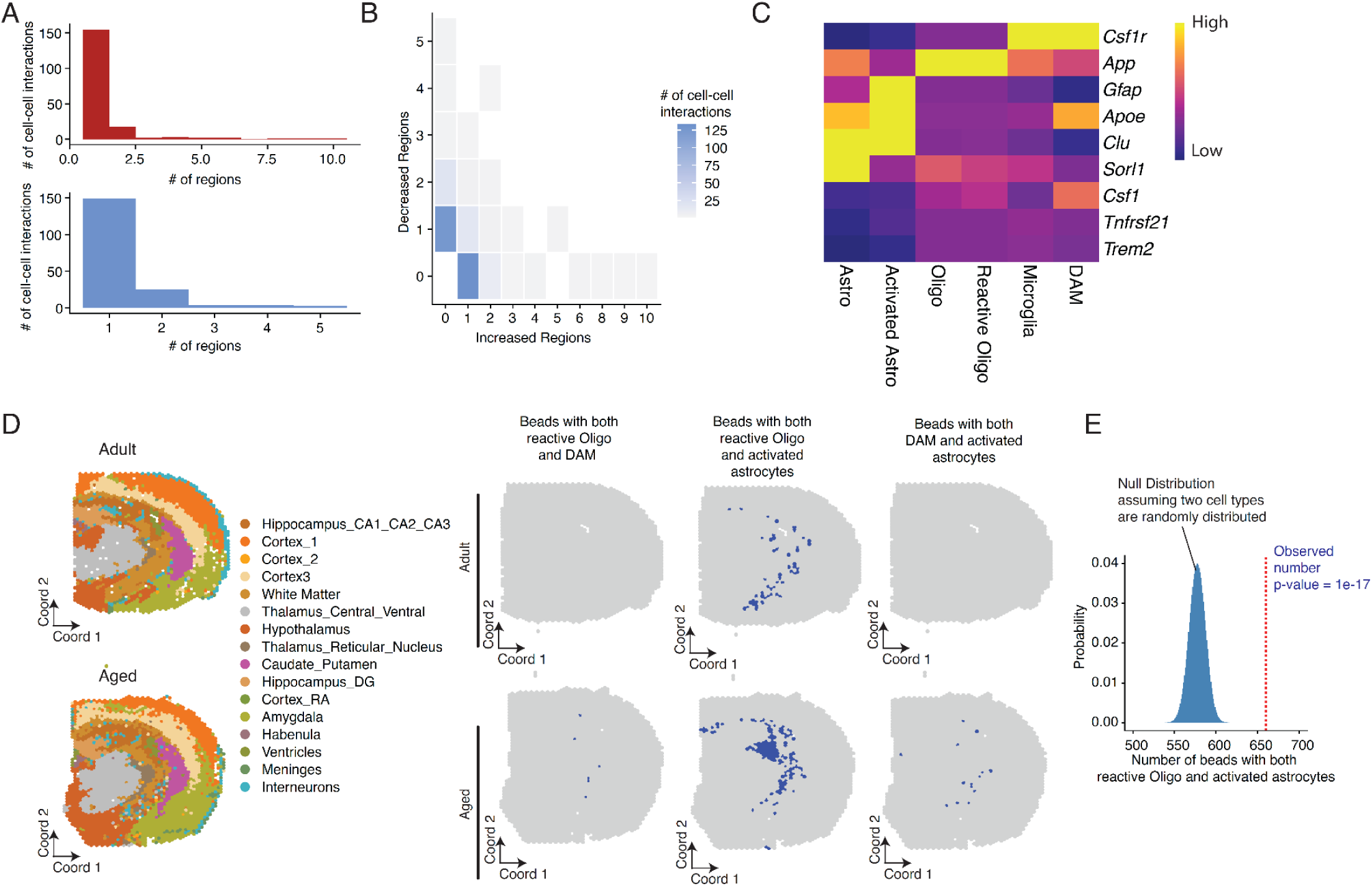
Region-specific alterations of cell-cell interactions in aging. (**A**) Histograms illustrating the number of cell-cell interactions that significantly increased (Top) or decreased (Bottom) across different brain regions. The x-axis represents the number of brain regions, while the y-axis shows the count of cell-cell interactions. (**B**) Heatmap displays the number of brain regions where each cell-cell interaction either expands (x-axis) or depletes (y-axis) with aging. The color gradient from light to dark red indicates the number of cell-cell interactions, with darker shades representing higher numbers. (**C**) Heatmap showing the normalized gene expression in DAM, reactive oligodendrocytes, activated astrocytes, and the corresponding main cell types. (**D**) The left panel illustrates the spatial distribution of transcriptomes across various brain regions in the adult and aged brain, color-coded by region. The right series of panels display the distribution of beads indicating colocalization of reactive oligodendrocytes and DAM, colocalization of reactive oligodendrocytes and activated astrocytes, and colocalization of DAM and activated astrocytes, shown for both adult and aged samples. (**E**) Histogram illustrating the null distribution of beads with both reactive oligodendrocytes and activated astrocytes assuming random distribution, with a dashed line marking the observed number.

**Table S1. Oligo sequences applied in the *IRISeq* protocol.** The 96-well plates contain oligonucleotide sequences for bead barcoding, designated for receiver and sender beads. The oligonucleotide sheets for receiver beads include: Ligation_1_Receiver_bead, Ligation_SP, Ligation_2, Ligation_2_SP, Ligation_3, Ligation_3_SP, and Ligation_4_Receiver_bead. The oligonucleotide sequences for sender beads are similar to those for receiver beads, with two exceptions. Firstly, during the initial round of barcoding, TruSeq Read2 is used in Ligation_1, and its sequences are found in the Ligation_1_Receiver_bead sheet. Secondly, Ligation_4 for sender beads differs by containing PolyA sequences instead of PolyT, with these sequences listed in the Ligation_4_Sender sheet. The SP sequences for the first round of ligation are identical for both sender and receiver beads. SP sequences for the fourth round of ligations, and PN2 for receiver and sender beads are included in Supplementary Note 1.

**Table S2. Metadata for mouse individuals included in the study.** For each animal and tissue section, this table describes the individual ID in the (“ID”) column, the section IDs associated with each individual in the (“section”) column, the date of birth (“DOB”) for each animal, the (“Gender”), the date of brain tissue harvested in (“Date harvested”), and date of library preparation in (“Date of Reverse Transcription”).

**Table S3. Region-specific gene features for reconstructed brain regions.** The Wilcoxon Rank Sum test in Seurat was used to analyze region-specific gene features by comparing each annotated brain region’s receiver beads to those of all other regions(*58*) (“avg_log2FC”) represents the log2 fold change of the average expression between the two groups, with positive values indicating higher gene expression in the first group. (“pct.1”) indicates the percentage of beads where the gene is detected in the region of interest, while (“pct.2”) indicates the percentage in the rest of the brain regions. (“p_val_adj”) denotes the Bonferroni-corrected p-value (“p_val”) for all genes in the dataset. “Region” refers to the brain region annotations, and “gene” shows the gene annotations and their reported values for each specific brain region.

**Table S4. Region-specific cell types for reconstructed brain regions.** For each annotated brain region, it was subsetted and binarized summed counts of cell subtype deconvolution values (as described in methods) were aggregated and each subtype proportion across the aggregated counts is calculated and reported as a percentage in the table. A minimum of 3% cutoff was used to generate subtype contributions to each region for this table. (“Brain_region”) highlights the annotated brain regions, (“Cell_Subtype”) highlights the subtype ID, and (“Percentage_contribution”) highlights (“Cell_Subtype”) contribution to the specific (“Brain_region”).

**Table S5. Aging-associated differentially expressed genes across brain regions.** For each gene, the “max.condition” is the condition with the highest expression (“max.expr”). The “second.condition” is the condition with the second highest expression (“second.expr”). The “fold.change” is the fold change between the max expression and second max expression. The “qval” is the false detection rate (one-sided likelihood ratio test with adjustment for multiple comparisons) for the gene differential expression test across different brain regions.

**Table S6. Aging-associated differentially abundant cell types across brain regions.** For each cell state (“cell_name”), the “max.condition” is the condition with the highest fraction of cellular abundance, calculated as the fraction of cells multiplied by 1 million to yield the “max.TPM” (Total Part per Million). The “second.condition” is the condition with the second highest fraction of cellular abundance (“second.TPM”). The “fold.change” is the fold change between the max and second max fraction. The “qval” is the false detection rate (one-sided likelihood ratio test with adjustment for multiple comparisons) for the differential abundance test across different regions.

**Table S7. Aging-associated cell-cell interactions across brain regions.** This table presents the differential abundance analysis results of cell-cell interactions similar to the cellular abundance analysis in Table S6.

## Notes

### Competing Interest Statement

J.C., W.Z., A.A., and W.J. are inventors on pending patent applications related to IRISeq. Other authors declare no competing interests.

