## Supplementary Note 1 for "Optics-free Spatial Genomics for Mapping Mouse Brain Aging"

### IRISeq protocol

#### Abstract
Here, we introduce *IRISeq* (Imaging Reconstruction using Indexed Sequencing), a novel, scalable, and cost-effective method for spatial genomic analysis that operates solely through sequencing, without the need for predefined capture arrays or optical imaging. *IRISeq* employs similar principles as those of DNA Hi-C (*1*) and DNA microscopy (*2*), decoding the spatial locations of molecules by sequencing their interactions with nearby molecules. Specifically, barcoded gel beads were utilized to capture gene expression locally, with the global spatial positions of these beads decoded through their interaction signals with adjacent beads.

#### Protocol workflow
The optimized *IRISeq* protocol comprises several key steps: (i) **Bead Fabrication**: Two types of oligo-barcoded beads are prepared: 'receiver beads' coated with PolyT sequences to capture nearby cellular mRNA, and 'sender beads' with a photocleavable linker and a PolyA sequence. These barcoded beads are created through a split-pool ligation approach (*3*), such that each bead has its unique barcode. (ii) **Photocleavage and Oligo Capture**: Beads are evenly distributed on a glass slide. A UV device is then utilized to photocleave oligos from the sender beads diffuse and are captured by the receiver beads, mimicking the capture of tissue mRNA. The beads array is then frozen on dry ice to stabilize the beads array. (ii) **Tissue Transfer and mRNA Capture**: Frozen tissue sections are then transferred onto this array using cryosectioning. The mRNA from the tissue is then captured by the receiver beads through hybridization and tissue digestion. (iv) **Reverse Transcription and Sequencing**: Post tissue digestion, beads are collected for reverse transcription and PCR, followed by sequencing to obtain transcriptome data and bead connection details.

#### Required Equipment
- iFlow Touch™ Microfluidic Pump System (Precigenome)
- Cryostat (Leica CM3050S)
- Centrifuge (Eppendorf 5702 RH)
- 12-tube Magnetic Separation Rack (NEB, S1509S)
- PCR tube Magnetic Separation Rack (Amazon)
- Eppendorf Mastercycler
- Freezer (-20C, -80C) and Refrigerator (4C)
- Gel Imager
- Ice Buckets
- Microscope
- Multi-channel Pipettes (2-20μL, 20-200μL) (Rainin Instruments)
- NextSeq 500 Platform (Illumina)
- Pipettors
- 96-well Pipetting System
- Eppendorf ThermoMixer C (5382000023) OR Fisherbrand Nutating Mixer (88861043)

#### Primer Sequences used

| Name | Oligo sequence |
|---|---|
| TSO | AAGCAGTGGTATCAACGCAGAGTGAATrG+GrG |
| dN-SMRT | AAGCAGTGGTATCAACGCAGAGTGANNNGGNNNB |
| Truseq_Read1 | ACACTCTTTCCCTACACGACGCTCTTCCGATCT |
| SMRT primer | AAGCAGTGGTATCAACGCAGAGT |

| | |
|---|---|
| P5_Truseq_Read1 | AATGATACGGCGACCACCGAGATCTACACACACTCTTTCCCTACACGACGCTCTTCCGATCT |
| 488-polyT probe | /5Alex488N/TTTTTTTTTTTTTTTTTTTTTTTTTTTTTT |
| 488-polyA probe | /5Alex488N/AAAAAAAAAAAAAAAAAAAAAAAAA |
| 5phos_3blocked_ME | /5Phos/CTGTCTCTTATACACATCT/3ddC/ |
| Tn5 Nextera R1 | TCGTCGGCAGCGTCAGATGTGTATAAGAGACAG |
| Tn5 Nextera R2 | GTCTCGTGGGCTCGGAGATGTGTATAAGAGACAG |
| Ligation_4_SP | /5Phos/CTCGAA TAGG |
| PN1-Acrydite_Receiver_Bead | /5ACryd/ACTAACAATAAGCTCTATCGATGACCTAATACGACTCACTATAGGGACACACTCTTTCCCTAC |
| PN1-Acrydite_Sender_Bead | /5ACryd/TTT/iSpPC/ACTAACAATAAGCTCTATCGATGACCTAATACGACTCACTATAGGGACGTGACTGGAGTTCAGAC |
| PN1-AmMC6_Receiver_Bead | /5AmMC6/ACTAACAATAAGCTCTATCGATGACCTAATACGACTCACTATAGGGACACACTCTTTCCCTAC |
| PN2_Receiver_Bead | /5Phos/ AGATCGGAAGAGCGTCGTGTAGGGAAAGAGTGT GTCCCTATAGTGAGTCGTATTAGGTCATCGATAGAG |
| PN2_Sender_Bead | /5Phos/ AGATCGGAAGAGCACACGTCTGAACTCCAGTCAC GTCCCTATAGTGAGTCGTATTAGGTCATCGATAGAG |

Barcoding primer sequences are attached in supplementary table 1. Bead barcoding split-pool barcoding primers are ordered from IDT with standard desalting. Primers are resuspended in water at 200 µM and normalized yield to the maximum available. Split primers are ordered at a larger scale due to their shorter length. 96 Well plate primers are ordered non-phosphorylated as in-house phosphorylation strategy is described below. All sequences are in supplementary table 1: Indexed P7 Truseq Read 2 primer sequences are included in sheet “Trueseq_Read_2_Indexed”. Indexed Nextera P7 Read 2 primer sequences are included in sheet “Nextera_Read_2_Indexed”.

**Materials used**

| Reagent | Vendor | Catalog |
|---|---|---|
| Acrylamide/Bis-acrylamide, 40% solution | Sigma | A9926-100ML |
| ACRYLAMIDE SOLUTION 40% 1L | Stockroom | 165000 |
| TEMED | Invitrogen | 17919 |

| | | |
|---|---|---|
| Ammonium persulfate | Sigma | A3678-25G |
| 20× SSC | Invitrogen | AM9763 |
| Tween 20 | Sigma | P9416-50ML |
| 1H,1H,2H,2H-Perfluoro-1-octanol | Sigma | 370533-25G |
| Maxima H minus Reverse transcriptase | Invitrogen | EP0753 |
| SUPERase• In™ RNase Inhibitor (20 U/µL) | Invitrogen | AM2696 |
| NEBNext® High-Fidelity 2X PCR Master Mix | NEB | M0541L |
| T4 DNA Ligase | NEB | M0202M |
| Fisherbrand™ Superfrost™ Plus Microscope Slides | Fisher Scientific | 12-550-15 |
| Single Emulsion Droplet Generator Chip - 4Device - Luer | Precigenome | 537 |
| Microfluidic Reservoir Kit for 15 mL Tube | Precigenome | N/A |
| Luer Lock Fittings Kit for Pressure Controller Connection, 4set/PK | Precigenome | N/A |
| Flangeles Fittings 1/4-28 to 1/16"OD, Nut and Ferrule, 10/pack | Precigenome | N/A |
| QX200™ Droplet Generation Oil for EvaGreen | Bio-Rad | 1864006 |
| Terra™ PCR Direct Polymerase Mix | Takara | 639271 |
| E-Gel™ EX Agarose Gels, 2% | Invitrogen | G402002 |
| Sodium hydroxide solution | Sigma | 72068-100ML |
| EDTA, 0.5 M, pH 8.0, Molecular Biology Grade, DEPC-Treated | Sigma | 324506-100ML |
| AMPure XP | Beckman Coulter | A63882 |
| E-Gel™ 50 bp DNA Ladder | Invitrogen | 10488099 |
| Mineral oil | Sigma | M5904-500ML |
| UltraPure™ 1M Tris-HCl, pH 8.0 | Invitrogen | 15568025 |

| | | |
|---|---|---|
| NaCl (5 M), RNase-free | Invitrogen | AM9759 |
| KCl (2 M), RNase-free | Invitrogen | AM9640G |
| T4 Polynucleotide Kinase | NEB | M0201L |
| Molecular Grade Water (nuclease-free) 1L | Coring | 46-000-CM |
| Recombinant albumin | NEB | B9200S |
| Buffer EB (250ml) | Qiagen | 19086 |
| DNA Binding Buffer | Zymo Research | D4004-1-L |
| $MgCl_2$ (1 M) | Invitrogen | AM9530G |
| Reagent Reservoirs 5mL | Genesee | 28-114 |
| Reagent Reservoirs 25mL | Thermofisher | 8093-11 |
| Exonuclease I (E. coli) | NEB | M0293L |
| Proteinase K | Thermofisher | 25530049 |
| Hemocytometer | VWR | 102966-632 |
| Brij™-35, 30% Solution | Thermofisher | 20150 |
| Zymoclean Gel DNA Recovery Kit (capped) | Zymo research | D4007 |
| dNTP | Thermofisher | R0193 |
| Klenow | NEB | M0212L |
| Triton X-100 | Sigma | T9284 |
| 20um filter | Pluriselect | 43-10020-60 |
| 100um filter | MACS | 130-110-917 |
| 1M Tris-HCl (pH 7.5) | Thermofisher | 15567027 |
| TE buffer 20X | Thermofisher | T11493 |

| | | |
|---|---|---|
| SDS, 10% Solution, RNase-free | Thermofisher | AM9822 |
| Ethanol | Stockroom | |
| Qubit dsDNA HS kit | Thermofisher | Q10212 |
| Qubit tubes | Thermofisher | Q32856 |
| Falcon Tubes, 15 ml | Stockroom | |
| Falcon Tubes, 50 ml | Stockroom | |
| DNA LoBind Tube 1.5 ml, PCR clean | Eppendorf | 22431021 |
| 96-well PCR Plate | geneseesci | 24-302 |
| 0.2mL 8-Strip Tubes with Individual Caps (PCR Tubes) | Geneseesci | 27-125U |
| eXTReme FoilSeal Film | Geneseesci | 12-156 |
| oYo-Link ® LED PX Device | alpha thera | AT8001 |
| eXTReme Clear Sealing Film | Southern labware | XTR-100 |
| Pipette Tips RT LTS 20uL FL 960A/10 | Rainin | 30389226 |
| Pipette Tips RT LTS 200uL F 960/10 | Rainin | 30389239 |
| Pipette Tips RT LTS 200uL FLW 960A/10 | Rainin | 30389241 |
| 10X genomic chamber | 10X genomics | |
| BcMag™ Quick oligo-DNA conjugation kit-5um | BioClone | CA-101 |
| PAMAM dendrimer, ethylenediamine core, generation 4.0 solution | Sigma | 412449-2.5G |
| DSS (disuccinimidyl suberate) | Thermofisher | 21655 |
| UltraPure™ Glycerol | Thermo Fisher | 15514011 |
| DMF: N,N-Dimethylformamide | Fisher Scientific | AC327175000 |
| Tn5 | UC Berkley | |

**Buffer Preparation**

1. Tris-buffered saline-EDTA-Triton buffer (TBSET) - storage in 4C for up to 6 months.

| | Stock | Final | Volume /ml |
|---|---|---|---|
| H2O | - | - | 465.6 |
| Tris-HCL pH8.0 | 1M | 10mM | 5.0 |
| NaCl | 5M | 137mM | 13.7 |
| EDTA pH8.0 | 0.5M | 10mM | 10 |
| Triton X-100 | 10% | 0.1%(v/v) | 5.0 |
| KCL | 2M | 2.7mM | 0.7 |
| | | | 500.0 |

2. Denature Butter -made fresh each time

| | Stock | Final | Volume /ml |
|---|---|---|---|
| H2O | - | - | 41.6 |
| NaOH | 1N | 150.00mN | 7.6 |
| Brij-35 | 30%(w/w) | 0.5%(w/v) | 0.8 |
| | | | 50.0 |

3. Tris-EDTA-Tween buffer (TET) - storage in 4C for up to 6 months.

| | Stock | Final | Volume /ml |
|---|---|---|---|
| Tris-HCL pH8.0 | 1M | 10mM | 5.0 |
| EDTA pH8.0 | 0.5M | 10mM | 10.0 |
| Tween 20 | 10%(v/v) | 0.1%(v/v) | 5.0 |
| | | | 500.0 |

4. Low salt buffer (LS) - storage in 4C for up to 6 months.

| | Stock | Final | Volume /ml |
|---|---|---|---|
| H2O | - | - | 488.9 |
| NaCl | 5M | 10mM | 1.0 |

| | | | |
|---|---|---|---|
| Tris-HCL pH8.0 | 1M | 10mM | 5.0 |
| EDTA pH8.0 | 0.5M | 10mM | 0.1 |
| Tween 20 | 10%(v/v) | 0.1%(v/v) | 5.0 |
| | | | 500.0 |

5. Pre ligation buffer (PL) - storage in 4C for up to 6 months.

| | Stock | Final | Volume /ml |
|---|---|---|---|
| H2O | - | - | 486.5 |
| NaCl | 5M | 30mM | 3.0 |
| Tris-HCL pH7.5 | 1M | 10mM | 5.0 |
| Tween20 | 10% | 0.1%(v/v) | 5.0 |
| Mgcl2 | 1M | 1mM | 0.5 |
| | | | 500.0 |

6. Tissue lysis buffer - storage at room temperature for up to one year.

| Reagent | Quantity (for 50 mL) | Final concentration |
|---|---|---|
| Tris-HCl (1 M, pH 8.0-8.5) | 5mL | 100 mM |
| NaCl (5 M) | 2ml | 200 mM |
| SDS (20% [w/v] in sterile H2O) | 5mL | 2% (w/v) |
| EDTA (0.5 M) | 500ul | 5 mM |
| H2O | 36.5 mL | |

7. Hybridization Buffer: 190ul 6X SSC buffer + 10ul RNAse inhibitor - made fresh each time.
8. TE-TW: 2.5ml 20X TE buffer + 50ul Tween 20 (10% solution, 0.01% final concentration) + 47.5ml ultrapure water. - storage at room temperature for up to 3 months.
9. 6X SSC (50ml): 15ml 20X SSC + 35ml ultrapure water. - storage at room temperature for up to 3 months.
10. 0.1N NaOH: 10ul 10M NaOH + 990ul H20. - made fresh each time.
11. 2D Buffer - 2x Tagmentation Buffer

1M Tris HCl (pH 7.5): 4mL

1M $MgCl_2$: 2mL DMF: 40mL
$H_2O$: add to 200mL (~154mL)
Filter buffer
Aliquot the solution into 15mL or 1.5mL tubes and store in -20C.

**Bead Fabrication**

Bead fabrication protocol is adapted from (3), with modifications noted in below protocol.
1. Prepare 10 ml of droplet generation oil with 0.6% v/v TEMED. Mix well in a 15 ml reservoir tube. (Note that the oil volume depends on the flow rates used and may require adjustments for different drop makers.)
2. Prepare the following three solutions:
*Note:* PN1 primers are diluted in water.

**Solution 1: Receiver Bead (PolyT)**

| Component | stock | | Volume/ul |
|---|---|---|---|
| Acrylamide/Bis-acrylamide, 40% solution 19:1 | 40% | w/v % | 100ul |
| Acrylamide 40% | 40% | | 100ul |
| Ammonium persulfate (APS) | 30% | w/v% | 15ul |
| TBSET | 100% | | 35ul |
| Tris-HCl pH8.0 | 1000 | mM | 50ul |
| PN1-Acrydite_Sender_Bead | 500 | uM | 200ul |
| Total | | | 0.5ml |

**Solution 2: Sender Bead (PolyA)**

| Component | stock | | Volume/ul |
|---|---|---|---|
| Acrylamide/Bis-acrylamide, 40% solution 19:1 | 40% | w/v % | 100ul |
| Acrylamide 40% | 40% | | 100ul |
| Ammonium persulfate (APS) | 30% | w/v% | 15ul |
| TBSET | 100% | | 35ul |
| Tris-HCl pH8.0 | 1000 | mM | 50ul |
| PN1-Acrydite_Receiver_Bead | 200 | uM | 200ul |
| Total | | | 0.5ml |

**Solution 3: Oil**

| Component | stock | final | | Volume/ul |
|---|---|---|---|---|
| TEMED | 100% | 0.6 % | v/v | 6ul |
| droplet generation oil | 100% | | | 994ul |
| Total | | | | 1000ul |

3. Load the reagents into the microfluidic reservoir tubes.
4. Note that there are two ports available on the reservoir cap: One port is the Luer lock connector, which connects the pressure source from the PreciGenome microfluidic controller. The other port is for reagent tubing, one side of which dips into the reagent, and the other side connects to the inlet of a microfluidic chip.
5. Connect the gas tubing to the reagent reservoir and the pressure controller integrated pressure pumps inside. Connect the reagent tubing to the reagent reservoir and the microfluidic chip. Ensure that the connections and fittings are airtight.
6. Set up the pressure controller appropriately for your experiment (Oil pressure ~8.2, Solution pressure ~5.8) . Start running the controller by simply clicking the "Run" button on the touch screen.
7. When the controller provides positive pressure, the reagents in the tubes are pushed to flow through the reagent tubing into microfluidic channels and then flow out to the waste.
8. Aim for a droplet diameter of 30 - 40 µm. Before collecting the droplets into the collection tube, take 5 µl of the droplets to a hemocytometer to check their size. If the size is in the desired range, collect the droplets into the first collection tube. Once a tube is full, close the lid and move it to an incubator set at 50°C for 1 hour or incubate the tube at room temperature overnight. *Note:* Overnight incubation at room temperature (RT) works equally well.

**Bead Cleanup**
**1. Centrifugation and Oil Removal:**
(1) Centrifuge the collection tubes at 100 g for 30 seconds.
(2) Use a 200 µl pipette tip to remove the droplet generation oil from the bottom of the tubes. Repeat this step multiple times to ensure thorough removal.
(3) Overlay the beads with 1 ml of TBSET and centrifuge at 3000 g for 30 seconds.
(4) Remove any leftover droplet generation oil and the mineral oil overlay, along with TBSET, by pipetting or aspiration.
(5) Repeat the TBSET washing step once more.

**2. PFO Treatment and Centrifugation:**
(1) Add 600 µl TBSET and 20 µl of 100% (v/v) 1H,1H,2H,2H-Perfluoro-1-octanol to each tube.
(2) Briefly vortex at full power.
(3) Centrifuge at 1500 g for 30 seconds.
(4) Use a pipette tip to remove the PFO mix from the bottom, followed by the mineral oil emulsion and TBSET from the top, without disturbing the beads.

**3. Bead Collection and Washing:**
(1) Place a 30 µm strainer on a 50 ml falcon tube and combine the beads from all tubes by transferring the bead emulsions to the strainer.
(2) Flush the strainer with TBSET until only bead precipitates remain on top.
(3) Transfer the strainer with beads precipitate on top to the top of a new tube, and flip over the strainer and flush the beads into the tube with TBSET.

**4. Centrifugation, Resuspension, and Filtering:**

(1) Centrifuge the tube at 1500 g for 3 minutes.
(2) Remove the supernatant and resuspend the pellet with 10 ml of TET.
(3) Transfer the bead suspension to a 15 ml tube.
(4) Filter the beads using a 100 μm filter to remove large beads.
(5) Wash the beads three times by centrifugation at 1500 g for 2 minutes, followed by resuspension with 10 ml TET.

*Stopping Point*: At this stage, the beads can be stored at 4°C for three months.

**For high resolution IRISeq only: bead Fabrication of 5 um sender bead (PolyT) with magnetic beads**
1. Prepare and activate the carboxyl groups on the 5 μm magnetic beads following the manufacturer's protocol (BcMag™ Quick oligo-DNA conjugation kit).
2. Prepare a dendrimer solution by mixing 500 μL of dendrimer solutions and 500 ul of manufacturer conjugation buffer, and incubate overnight on a rotator.
3. After incubation, wash the beads according to the manufacturer's protocol. Perform multiple rounds of washing with 1X PBS.
4. Bead Conjugation with PN1 oligonucleotide:
(1) Prepare 100 μL of 25 mM DSS solution fresh and place on ice.
(2) Mix 100 μL of 200 μM PN1-AmMC6_Receiver_Bead primer with 500 μL of 1X PBS.
(3) Add the fresh 20 μL DSS solution to the oligonucleotide mixture.
(4) Incubate the beads with the PN1 primer mixture with DSS on a rotator for 1 hour at room temperature.
(5) Inhibit the reaction by washing the beads with 1 mL of 1M Tris-HCl (pH 7.5) and then performing multiple rounds of washing with 1X PBS.

*Store the conjugated beads at 4°C in water and/or PBS for several weeks until barcoding is performed.
*During split-pool barcoding, beads are washed on a magnetic separator instead of spinning.

**Beads Barcoding**
1. Calculate the volumes required based on the bead volume used (1 ml or 5 ml). Ensure that the PCR plate can hold the total volume.
2. Thaw all barcode plates with the added splints mixed in a 1:1 ratio, for a final concentration of 100 uM when combined with splits.
*Note:* All split-pool ligation sequences are found in supplementary table 1.

*Phosphorylation of Primers:*
1. Prepare the T4 polynucleotide kinase (PNK) master mix for each plate.

| | | | | |
|---|---|---|---|---|
| T4 lig buffer | 10 | 1.00 | x | 600ul |
| H2O | - | - | - | 3550ul |
| BSA | 20 | 0.06 | mg/ml | 18ul |
| T4 PNK | 10 | 0.05 | U/ul | 32ul |

2. Add 40 μl master mix to each well of a PCR plate.
3. Add 20 μl from each well of the Ligation_1_Receiver primers mixed with Ligation_1_SP (final 100 uM concentration) to the corresponding well of the PCR plate using a multi-channel pipette. Mix by pipetting up and down. Similar reaction is performed for sender beads barcoding, with an exception to use Ligation_1_Sender beads primer plate.

4. Seal the plate with aluminum foil and incubate at 37°C for 30 minutes.
5. Heat-inactivate the reaction at 65°C for 20 minutes.

*Split-Pool Protocol, 1st Round:*
1. While primers are phosphorylating, wash beads 3 times with 10 ml PL buffer.
2. Resuspend the beads according to the following table and heat to 75°C for 2 minutes.
3. Let the mix cool down to room temperature slowly to anneal the beads and PN2 primer.

| | | | | |
|---|---|---|---|---|
| Beads | - | - | - | 500ul |
| T4 ligase buffer | 10 | 1 | X | 100ul |
| PN2_Receiver_Bead for receiver beads or PN2_Sender_Bead for sender beads | 100 | 3.75 | uM | 75ul |
| H2O | - | - | - | 325ul |

4. Wash the beads with PL buffer 3 times to remove extra single PN2.
5. Prepare the mixture including beads and T4 ligase buffer.

| | | | | |
|---|---|---|---|---|
| Beads | - | - | - | 500ul |
| T4 ligase buffer | 10 | 1 | X | 100ul |
| H2O | - | - | - | 400ul |

*Ligation:*
1. Once primer phosphorylation is complete, remove the foil and distribute 20 µl beads to each well.
2. Store a 5 µl bead aliquot in a PCR tube, labeled "round 0" for quality control.
3. Prepare the T4 ligase master mix in 5 ml tubes and distribute 40 µl to each well. Mix by pipetting up and down. Use a multi-channel pipette and a reagent reservoir. Replace tips after each pipette cycle to avoid cross-contamination.

| | | | | |
|---|---|---|---|---|
| T4 ligase buffer | 10 | 1 | x | 420ul |
| H2O | - | - | - | 3760ul |
| T4 ligase | 2000 | 1.91 | U/ul | 20ul |

4. Seal the plate and incubate at room temperature for 1 hour. Heat-inactivate the reaction at 65°C for 10 minutes and cool to room temperature.

*Washing:*
1. Use a multi-pipette to collect beads into a reagent reservoir filled with 3 ml TET. No need to replace tips in this step.

2. Collect beads from reagent reservoir to 15ml tubes. Centrifuge at 1000 g for 3 minutes and remove the supernatant.
3. Wash five times in 10 ml TET.

*Note: 5 um receiver beads are washed on the magnetic separator, and not by spinning.*

*Split-Pool Protocol, 2nd, 3rd, and 4th Round:*
1. Proceed as for the first round barcoding using barcode plate 2, barcode plate 3 and barcode plate 4.
2. No need to add more PN2 for either receiver or sender beads since the barcode fragments are already double-stranded. Replace the PN2 volume with water.
3. Always store a 5 µl bead aliquot in a PCR tube, labeled with the corresponding cycle number for quality control.

*Note*: barcode plate 1 and 4 for receiver and sender beads are different.

*Denaturing and Quenching:*
1. Pellet beads at 2000 g for 3 minutes, remove supernatant.
2. Wash the beads 3 times with 10 ml denaturing buffer to make the conjugated primer single-stranded.
3. After the last wash, remove the supernatant and wash 3 times with 10 ml LS buffer to quench the denaturing buffer.

*Note: 5 um receiver beads are washed on the magnetic separator, and not by spinning.*

**Quality Control of Beads by Fluorescence Imaging**
1. Resuspend the quality control beads in 20ul TET buffer and transfer 5µl aliquots to PCR tubes.
3. Add 25 µl TET to each tube.
4. Add 1 µl of fluorescence probe (100 µM stock) to the tube. Use 488-polyT probe for sender bead, and 488-PolyA for receiver bead.
5. Vortex and incubate the beads with probes for 30 minutes at room temperature in the dark.
6. Wash the beads 3 times with 1 ml of TET.
7. Image beads under a fluorescence microscope.

**Note: Sequences for each ligation barcode are provided in Supplementary Table 1.** Before the barcoding process, mix 200 uM of each Ligation and Ligation Splints for a final concentration of 100 uM. The mixed primer plate can be stored at -20 C for up to a year.

**Receiver Bead ligation plate combinations**

| | Ligation | Ligation Splint |
|---|---|---|
| 1st round ligation-Plate 1 | **Ligation_1_Receiver_bead** | **Ligation_1_SP** |
| 2nd round ligation-Plate 2 | **Ligation_2** | **Ligation_2_SP** |
| 3rd round ligation-Plate 3 | **Ligation_3** | **Ligation_3_SP** |
| 4th round ligation-Plate 4 | **Ligation_4_Receiver** | **Ligation_4_SP** |

Note: PN2_Receiver_Bead for first round of ligation

**Sender Bead ligation plate combinations**

| | Ligation | Ligation Splint |
|---|---|---|
| 1st round ligation-Plate 1 | **Ligation_1_Sender_bead** | **Ligation_1_SP** |

| | | |
|---|---|---|
| 2nd round ligation-Plate 2 | **Ligation_2** | **Ligation_2_SP** |
| 3rd round ligation-Plate 3 | **Ligation_3** | **Ligation_3_SP** |
| 4th round ligation-Plate-4 | **Ligation_4_Sender** | **Ligation_4_SP** |

Note: PN2_Sender_Bead for first round of ligation

**Library Preparation and Sequencing**

Library preparation is modified from Slide-seq V2(4). Specifically, we have developed three versions of protocol for tissue RNA capture with different beads: 1) IRISeq with Frozen bead array: this is the most optimized version of IRISeq compatible for profiling both small and large bead array; 2) IRISeq with non-frozen bead array: this is a developmental version of the protocol with slightly lower efficiency and purity. 3) IRISeq with 5um magnetic beads: this is an alternative version of IRISeq for spatial transcriptome profiling at high resolution.

**Tissue Preparation:**

1. Mice were anesthetized with CO2 and decapitated.
2. The brain was rapidly dissected, frozen on crushed dry ice, and stored at −80 °C until cryosectioning.
3. Warm fresh frozen tissue to -20 C in a cryostat for 20 minutes.
4. The tissue was mounted onto a cutting block with OCT and sliced at 10 μm thickness for RNA capture.

**IRIS-seq with the frozen bead array (Optimized protocol for both small and large areas profiling):**

1. Cut a 0.6 cm x 0.6 cm (or 1.5cm x 1.5cm for large array) square-shaped area in a 96-well plate plastic tape. Tape the cut area onto a glass slide to define the square shape of the bead layer.
2. Prepare a 15ul mixture of receiver beads and sender beads in a 3:1 ratio in hybridization buffer (Prepare 50 ul 3:1 mixture for 1.5 cm x 1.5 cm array size). Place 5ul bead mixture on the glass slide in the taped square-shaped area, making sure all area is covered with gel beads (Place ~30 ul of beads mix for 1.5 cm x 1.5 cm array, and add more beads accordingly to fill the square). Add 1 μl of Proteinase K to the bead mixture. Homogenize the beads using a pipette tip to facilitate the beads to form a monolayer. (Add 5 ul for Proteinase K for 1.5 X 1.5 cm sized array)
3. Incubate the beads array at room temperature for 1-2 minutes to allow the surface to form a monolayer. Take care to make sure the array is not too dry as this reduces barcode diffusion. Then, place the beads array in a photocleavage chamber for 2 minutes.
4. After photocleavage, immediately place the beads array on dry ice to freeze and stabilize the array. Transfer the tissue section onto the slide. Place the beads array with the tissue on dry ice again and transfer it to a 37°C humid chamber for 20 minutes for tissue digestion and RNA capture.
5. Collect the beads in a 15 ml tube using a funnel. Wash the glass array in the funnel with 6XSCC to collect the beads. Centrifuge the tube at 2000 g for 5 minutes. Collect the supernatant and wash the beads twice more with approximately 5 ml of 6XSCC.

* IRIS-seq with the non-frozen bead array (Developmental version of the protocol for profiling 0.6cm x 0.6cm sections):

1. Cut a 0.6 cm x 0.6 cm (or 1.5cm x 1.5cm for large array) square-shaped area in a 96-well plate plastic tape. Tape the cut area onto a glass slide to define the square shape of the bead layer.
2. Prepare a 15ul mixture of receiver beads and sender beads in a 3:1 ratio in hybridization buffer. Place 5ul bead mixture on the glass slide in the taped square-shaped area, making sure all area is covered with gel beads
3. When the beads array was slightly dry and no liquid layer was observed, the slide was placed on the cutting stage in the cryostat and cooled for ~10 seconds, and then 10 μm tissue was transferred to the bead array.
4. The tissue was melted onto the slide by moving the glass slide off the stage and placing a finger on the bottom side of the glass.
5. Very carefully, the slide was removed from the cryostat and placed into a humidity glass slide chamber with liquid at bottom of chamber to keep tissue humid and not dry.
6. The array in the chambered slide with tissue on top was incubated at room temperature for 13 minutes.
7. Photocleavage of the beads array was performed for 2 minutes under UV light using (oYo-Link LED PX Device), an ice chilled metal plate is used to prevent tissue from becoming too dry during photocleaving.
8. A 10x genomic gasket chamber, with a 20 μm filter between the gasket and the beads array, was placed to stabilize tissue on the beads during reverse transcription.

* For high-resolution IRISeq with 5 um receiver beads, 10 um tissue sections are cut and placed on a glass slide, and then a mixture of 5 um receiver beads with 50 um sender beads (~80,000 5 um beads, with ~8,000 50 um sender beads, add more beads accordingly depending on tissue size to cover whole tissue area) are mixed in the hybridization buffer. An aliquot of the mixture was taken and monodispersed on the tissue. Hybridization, photocleaving, and gasket assembly are similar to the description above.

**Reverse Transcription**

The sample library was then prepared as below. (The remaining tissue was re-deposited at -80 C and stored for processing at a later date). The reaction volume below is for .6 X .6 cm sized array tissue capture. For 1.5 X 1.5 cm array tissue capture, reaction conditions are multiplied by 5X. Beads are separated into 5 tubes following tissue RNA capture, with each tube counted as a .6 X .6 cm reaction (same reaction mixture concentrations described below), and all samples are merged before library preparation for sequencing.

1. Prepare the RT reaction buffer (can be prepared in advance without RTase):
    - 120 µL H2O
    - 40 µL Maxima 5x RT Buffer
    - 20 µL 10 mM dNTPs
    - 5 µL RNase Inhibitor (ThermoFisher)
    - 5 µL 100 uM Template Switch Oligo (TSO)
    - 10 µL Maxima H- RTase
    - Total volume 200 µl
2. Add 200 µl of RT mix to the beads (very gently to avoid bubbles on the filter for hybridization-based RNA capture). Reverse transcription is performed in the tube with beads for IRISeq with the frozen bead array or on the beads array for IRIS-seq with the non-frozen bead array.
3. Incubate the reaction at 50°C for 90 minutes (use PCR machine glass slide adapter for IRISeq with the non-frozen bead array).

**Tissue Lysis (IRISeq with non-frozen bead array only)**

1. Prepare a working solution for tissue digestion by adding proteinase K to the tissue-clearing buffer stock solution at a 1:50 ratio:

Working Solution:

- 196 µL Tissue lysis buffer
- 4 µL Proteinase K enzyme

*Note:* The tissue lysis buffer (without proteinase K) can be made in advance and stored at room temperature for several months.

2. Add 200 µL of the tissue lysis mix to the beads in each tube, bringing the total volume to 400 µL.
3. Incubate the mixture at 37°C for 30 minutes.
4. Cut the filter and collect all beads as much as possible into a PCR tube.

*Note:* This is a stopping point. The beads can be stored at 4°C in TE-TW for overnight.

**Exonuclease I Treatment**

1. Prepare Exonuclease I mix recipe (200 µL):
    - 20 µL 10X Exo I Buffer
    - 170 µL H2O
    - 10 µL Exo I
2. Pellet and then wash the beads with 10 mM Tris-HCl, pH 7.5.
3. Pellet beads again and resuspend the beads in 200 µL Exonuclease I mix. Incubate the reaction at 37 °C for 50 minutes.
4. After Exonuclease I treatment is done, pellet the beads and remove supernatant. Wash beads twice with TE-TW.

5. Pellet beads once more and resuspend the beads in 200 µL 0.1 N NaOH for 5 minutes at room temperature. Note: Prepare 0.1 N NaOH immediately before use.
6. Quench NaOH with 200 µL TE-TW.
7. Pellet beads as before, wash once with TE-TW and once with 1x TE buffer.

**Second Strand Synthesis**
1. Prepare second strand synthesis mix per puck:
   - 133 µL ultrapure water
   - 40 µL Maxima 5x RT Buffer
   - 20 µL 10 mM dNTPs
   - 2 µL 1 mM dN-SMRT oligo
   - 5 µL Klenow Enzyme
     Total volume 200 µL
2. Pellet beads and resuspend the beads in 200 µL second strand synthesis mix. Incubate the reaction at 37 °C for 1 hour.

**cDNA Library Amplification**
1. Resuspend the beads in 200 µl the TE-TW buffer and transfer them to new DNA Lo-bind tubes.
2. Repeat the washing step with the TE-TW buffer for three times.
3. Pellet the beads once more.
4. Add 80 µl of 0.1 N NaOH to the bead pellets and incubate for 5 minutes.
5. Pellet the beads and collect the supernatant into a tube.
6. Quench the reaction with 13.5 µl of 1 M pH 7 Tris buffer.
7. Perform 1X Ampure beads purification and elute the cDNA in 30 µl EB buffer.
8. Prepare the following PCR mix per tube:
   - 30 µl Elution supernatant
   - 60 µl Ultrapure water
   - 100 µl Terra PCR Direct Buffer
   - 3 µl 100 µM P5_Truseq_ Read1 primer
   - 3 µl 100 µM SMRT PCR primer
   - 4 µl Terra Polymerase
   - Total volume 200 µl
9. Divide the total volume of each sample into two PCR tubes, each containing 100 µl (50%) of the total. Run the following PCR program:
   - 95 °C for 3 minutes
   - 4 cycles of:
     - 98 °C for 20 seconds
     - 65 °C for 45 seconds
     - 72 °C for 3 minutes
   - 9 cycles of:
     - 98 °C for 20 seconds
     - 67 °C for 20 seconds
     - 72 °C for 3 minutes
   - Then:
     - 72 °C for 5 minutes
     - 4 °C forever

**cDNA Library Purification**
1. Transfer 200 µL of PCR product into a 1.5ml tube.
2. Perform 0.6X Ampure beads purification and elute the cDNA in 22 µl water.
3. Normalize the library concentration to 3 ng/ul using water.

*Expected Result*: Your cDNA library should exhibit a relatively consistent and smooth appearance, with an average fragment size ranging from 1000 to 1500 bp as shown below in a representative gel image.

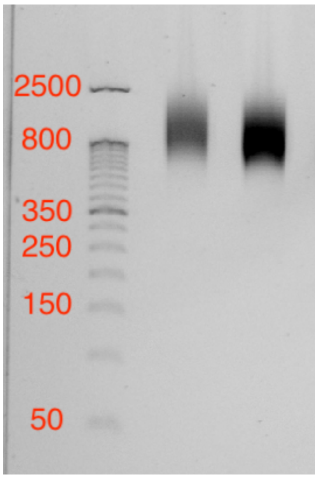


**Receiver bead-sender bead Connections Library: PCR and Gel Extraction**

1. Wash the beads with TE-TW twice, then with water, and suspend them in 27 µL of H2O.
2. Prepare a PCR mix containing:
    - Beads suspended in 27 µL of H2O
    - 1.5 µL of 10 uM Truseq_indexed_P7_Read2
    - 1.5 µL of 10 uM P5_Truseq_ Read1
    - 30 µL of NEBnext master mix
    - Total volume: 60 µL
3. Run the following PCR program:
    - Start
    - 72 °C for 3 minutes
    - 95 °C for 30 seconds
    - 12 cycles of:
        - 95 °C for 10 seconds
        - 55 °C for 10 seconds
        - 72 °C for 20 seconds
    - Then:
        - 72 °C for 3 minutes
        - 4 °C, hold
4. After PCR, take 3 µL to run on a gel. Then, cut the desired band and perform gel extraction according to the manufacturer's instructions using Zymoclean Gel DNA Recovery Kit. Bands for receiver-sender bead connections are highlighted below in a representative gel image.

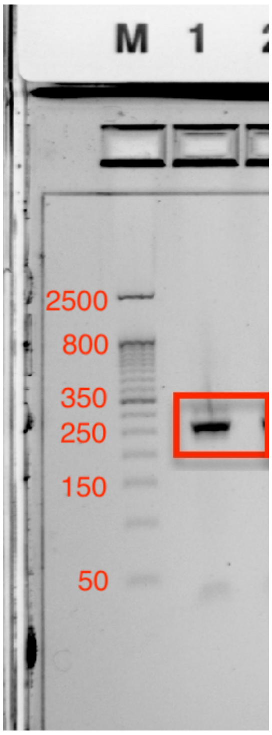


5. Finally, normalize the library concentration to 2 nM with water for sequencing.

**Tn5 loading.**

*Prepare buffer* -Dialysis buffer Recipe: 50 mM Tris-HCl pH 7.5, 800 mM NaCl, 0.2 mM EDTA, 10% glycerol

1. Prepare the Nextera-P7/P5 Tn5: Mix 100 μL of Tn5 dialysis buffer with 2 μL of 100 mM DTT.
2. Prepare the Oligos for Annealing for each end as shown below:

**For End 1**

| **Tn5_Nextera_R1** | Stock | Final | 1X | 20X |
|---|---|---|---|---|
| Tn5_Nextera_R1 | 100uM | 8uM | 0.4ul | 8ul |
| 5phos_blocked_ME | 100uM | 8uM | 0.4ul | 8ul |
| Tn5 dialysis (with 2mM DTT final) for each end | | | 1.7ul | 34ul |
| Total for each end | | | 2.5ul | 50ul(for each end: 8ul+8ul+34ul=50ul ) |

**For End 2**

| **Tn5_Nextera_R2** | Stock | Final | 1X | 20X |
|---|---|---|---|---|
| Tn5_Nextera_R2 | 100uM | 8uM | 0.4ul | 8ul |
| 5phos_blocked_ME | 100uM | 8uM | 0.4ul | 8ul |
| Tn5 dialysis/dilution buffer (with 2mM DTT final) for each end | | | 1.7ul | 34ul |

| | | | | |
|---|---|---|---|---|
| Total for each end | | | 2.5ul | 50ul(for each end: 8ul+8ul+34ul=50ul) |

3. Run the PCR program in a thermocycler to anneal the oligonucleotides:
(1) Start at 95°C for 5 minutes.
(2) Slowly cool down to 65°C at a rate of 0.1°C per second.
(3) Hold at 65°C for 5 minutes.
(4) Slowly cool down to 4°C at a rate of 0.1°C per second.
(5) Combine the two annealed oligo mixes to obtain 100 μL (50 μL * 2) of 16 μM annealed Tn5 oligo mix (8 μM for each end).

4. Prepare Glycerol Tn5 Stock (~50%):
(1) Thaw 30 μL of Tn5 extract on ice (original concentration: 4 mg/mL).
(2) Prepare 90 μL of diluted glycerol buffer (30 μL Dialysis buffer + 60 μL 100% ultrapure glycerol + 1.8 μL of 100 mM DTT), mix gently by pipetting until no separation of layers.
(3) Add the thawed Tn5 to the diluted glycerol buffer, and mix by gentle rotation (15rpm) at 4°C for 20 minutes.
(4) Gently mix the reaction several times by a P1000 pipette, and keep the tube on ice until Tn5 loading.

5. Load Nextera R1/R2-Tn5:
(1) Add 110 μL of prepared Glycerol Tn5 Stock stock to the combined annealed oligo mix tube and mix gently by pipetting for several times.
(2) Incubate the reaction at 25°C for 30 minutes with gentle shaking on the thermomixer at 300 rpm.
(3) Store at -20°C until needed.

**cDNA Library Tagmentation and PCR**

1. Preheat the PCR machine to 55°C.
2. Prepare the Tn5 reaction in a PCR tube:
    - 2.5 μl 2X TD buffer
    - 0.5 μl Tn5
    - 2 μl DNA (3 ng/ul working concentration)
    - Total volume: 5 μl
3. Incubate the PCR tube at 55°C for 5 minutes.
4. Add 5 μl DNA binding buffer to the PCR tube, and mix well.
5. Perform 1X Ampure beads purification and elute the cDNA in 9 μl Elution buffer.
6. Prepare the PCR mix:
    - 9 μl Eluted DNA from the previous step
    - 1 μl 10 uM P5_Truseq_ Read1primer
    - 1 μl 10 uM Nexterra_indexed_P7_Read2 primer
    - 11 μl 2X NEBnext Master Mix
    - Total volume: 22 μl
7. Run the following PCR program:
    - Start:
        72°C for 3 minutes
        95°C for 30 seconds
    - 13 cycles of:
        95°C for 10 seconds
        55°C for 10 seconds
        72°C for 30 seconds
    - Then:
        72°C for 3 minutes

4°C, hold

**Library Purification**

1. Add an additional 18 µl of H2O to the PCR mix, increasing the total volume to 40 µl.
2. Perform 0.8X Ampure beads purification and elute the cDNA in 12 µl water.
3. Normalize the library concentration to 2 nM for sequencing.

**Note: PCR Library Characteristics**

Your tagmented library should be fairly smooth, with an average bp size of 300-400bp. As shown below.

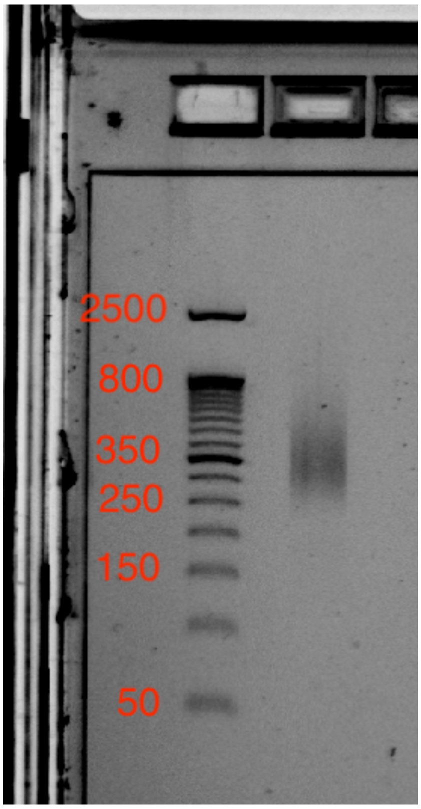
